# Essential dynamic interdependence of FtsZ and SepF for Z-ring and septum formation in *Corynebacterium glutamicum*

**DOI:** 10.1101/732925

**Authors:** Adrià Sogues, Mariano Martinez, Quentin Gaday, Mathilde Ben-Assaya, Martin Graña, Alexis Voegele, Michael VanNieuwenhze, Patrick England, Ahmed Haouz, Alexandre Chenal, Sylvain Trépout, Rosario Duran, Anne Marie Wehenkel, Pedro Alzari

## Abstract

The mechanisms of Z-ring assembly and regulation in bacteria are poorly understood, particularly in non-model organisms. *Actinobacteria*, one of the largest bacterial phyla that includes the deadly human pathogen *Mycobacterium tuberculosis*, lack the canonical FtsZ-membrane anchors as well as all positive and negative Z-ring regulators described for *E. coli*. Here we investigate the physiological function of *Corynebacterium glutamicum* SepF, the only cell division-associated protein from *Actinobacteria* known to directly interact with the conserved C-terminal tail of FtsZ but whose actual mode of action in cytokinesis is yet to be elucidated. We used a mechanistic cell biology approach to unveil the essential interdependence of FtsZ and SepF required for the formation of a functional Z-ring in the actinobacterial model organism *C. glutamicum*. The crystal structure of the SepF-FtsZ complex reveals a hydrophobic FtsZ-binding pocket, which defines the SepF homodimer as the functional unit, and a reversible oligomerization interface regulated *via* an alpha helical switch. FtsZ filaments and lipid membranes have opposing effects on SepF polymerization, leading to a complex dynamic role of the protein at the division site, involving FtsZ bundling, Z-ring tethering and membrane reshaping activities that are needed for proper Z-ring assembly and function.

---

The prokaryotic tubulin homologue FtsZ is at the heart of bacterial cytokinesis. At the cell division site, FtsZ protofilaments form a highly dynamic, membrane-bound structure, the Z-ring, which serves as a scaffold for the recruitment of the extra-cytoplasmic cell wall biosynthetic machinery. Despite its discovery more than 25 years ago ^1^ the exact molecular mechanisms of Z-ring assembly and regulation remains enigmatic ^2^. In the best studied organisms such as *Escherichia coli* and *Bacillus subtilis*, the action of several auxiliary proteins is necessary to positively or negatively regulate Z-ring formation (EzrA, ZapA-D, ^3–7^), to tether the structure to the membrane (FtsA, ZipA, SepF^8–11^) and to ensure proper subcellular (midcell) localization (SlmA, Noc, MinC/MinD, ^12,13,14^). Most of these proteins exert their functions by binding directly to the highly conserved FtsZ C-terminal domain (FtsZ_CTD_) ^15^, which is separated from the core GTPase domain by an intrinsically disordered linker of variable length and sequence. Interestingly, except for SepF all the above positive and negative FtsZ regulators are missing in *Actinobacteria* ^16^, for which cell division mechanisms are largely unknown.

The *sepF* gene is found in the *dcw* (division and cell wall) cluster along with *ftsZ* and many essential genes for cell division ^17^. In *B. subtilis*, SepF is a non-essential membrane-binding protein that co-localizes with FtsZ at mid-cell and is required for correct septal morphology as part of the late divisome ^18,19^. In contrast to *B. subtilis*, *sepF* is an essential gene in *Mycobacterium smegmatis* ^20^ and in the cyanobacterium *Synechocystis* ^21^, both of which lack an identifiable homologue of *ftsA*. In *M. smegmatis* SepF localizes to the Z-ring in a FtsZ-dependent manner and has been shown to interact with the conserved C-terminal domain of FtsZ in yeast-two-hybrid assays ^20^. Like FtsA, SepF has self-associating properties ^22^ and thus appears as a likely candidate for FtsZ membrane tethering in *Actinobacteria*. However, the observed assembly of SepF into stable 50 nm diameter ring polymers (alone or by bundling FtsZ protofilaments) seems to lack the dynamic oligomerization properties that are a recurrent feature of divisome and Z-ring interactors ^23,24^. Indeed, increasing evidence suggests that membrane anchors are not just passive but active players of Z-ring dynamics and regulation. For instance, FtsA has a dual role both serving to tether FtsZ filament fragments to the membrane and exercising an antagonistic function on polymerization dynamics and directional assembly at mid-cell, a pre-requisite to robust proto-ring assembly and subsequent inwards growth of new cell wall ^10,24,25^. While in *B. subtilis* FtsA and SepF may have complementary functions ^11^, we asked what happens in species where only SepF is present as a major Z-ring membrane anchor. Here we provide mechanistic insights for the FtsZ-SepF interaction and its interdependency for Z-ring assembly and regulation in *C. glutamicum*. We show that SepF has a complex dynamic role at the division site and that the ternary interaction between SepF, FtsZ and the membrane, coupled to FtsZ polymerization dynamics, are all required for proper function and assembly.

## Results & Discussion

### The essential role of SepF in *C. glutamicum*

SepF from *Mycobacterium smegmatis* was shown to be essential for viability ^20^ and it was proposed that, since FtsA is absent in *Actinobacteria*, SepF would be the unique membrane anchor for FtsZ. However, an early study reported that *sepF* was not essential in *Corynebacterium glutamicum* ^26^, which would argue against a main membrane tethering role for SepF. Several attempts at deleting *sepF* from *C. glutamicum* using either homologous recombination or gene disruption failed, suggesting that this gene might indeed be essential for bacterial survival. To deplete *sepF* we designed a mutant strain (*P*_*ino*_-*sepF*) where the transcription of *sepF* was uncoupled from its physiological promoters by placing a transcriptional terminator just before the *sepF* gene, and by putting it under the control of the previously described *myo*-inositol repressible promoter (*P*_*ino*_) of the inositol phosphate synthase Ino1 gene ^27^. Downstream effects were not expected as *sepF* is the last gene to be transcribed in the *dcw* cluster in *C. glutamicum* ^28^. We observed a rapid depletion of SepF in the presence of 1% *myo*-inositol, while in its absence the SepF protein levels remained close to the level in the wild-type (WT) strain (Fig. 1a). The growth curves of the depleted *P*_*ino*_-*sepF* and WT strains followed a similar pattern during the first 6 hours, but after that point growth stopped for the depleted strain (Fig. 1b). When observed under the microscope a strong phenotype was seen from the first time point (t=3h), with elongated cells that eventually lysed (Figs. 1c, d). At later time points branching was also seen, which corresponds to the formation of new poles at misplaced sites over the lateral walls of the bacterial cell. This *sepF* depletion phenotype was rescued when the strain was complemented with a plasmid carrying an extra copy of *sepF* under the control of the *P*_*tet*_ promoter (Supplementary Fig. 1) thus demonstrating the essentiality of *sepF* in *C. glutamicum*.

**Figure 1:**
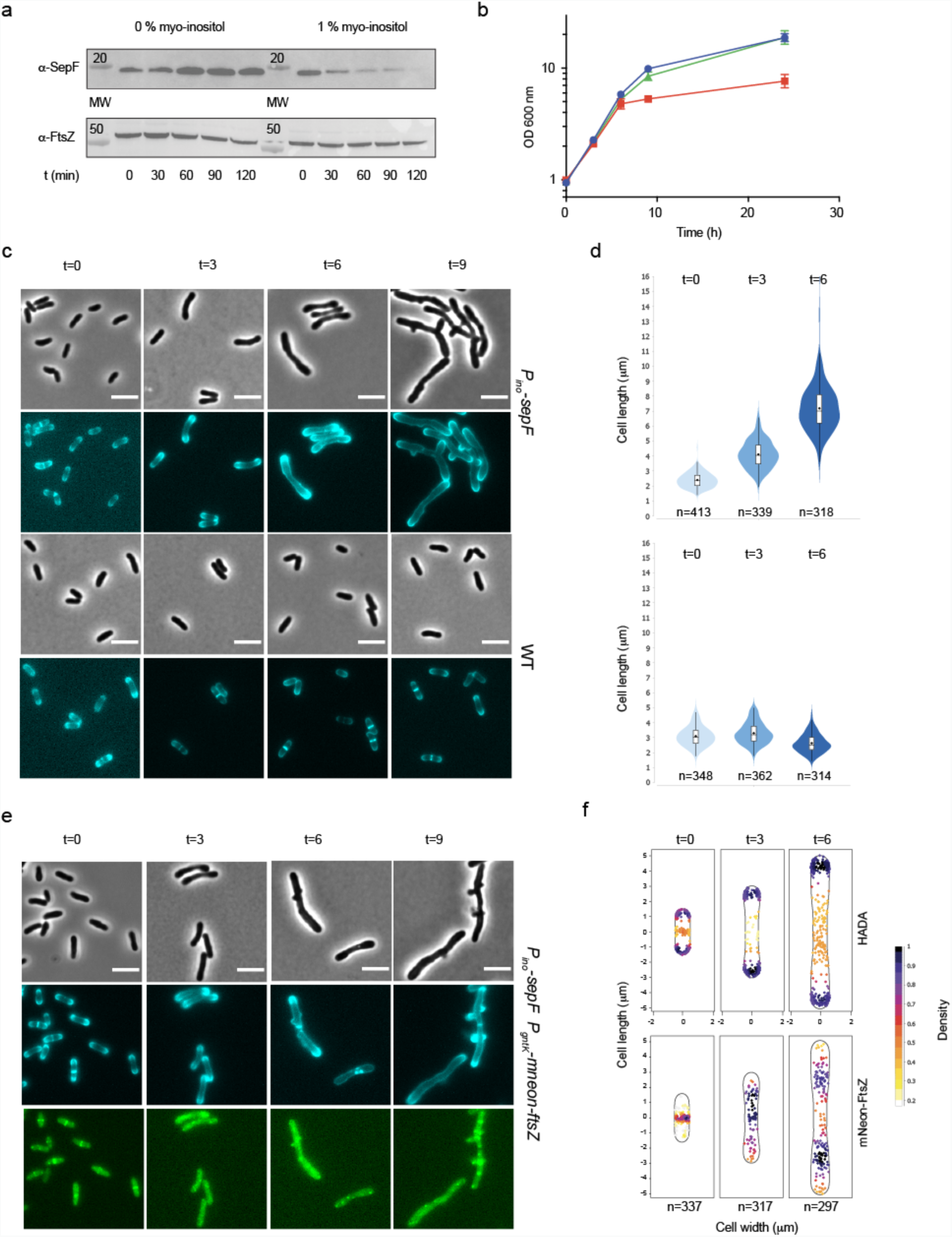
Phenotypic characterization of *P*_*ino*_-*sepF*. **a.** SepF depletion. Western blots of whole cell extracts from the *P*_*ino*_-*sepF* strain, in the absence (not depleted) or presence (SepF depleted) of 1% *myo*-inositol during 2 hours. SepF and FtsZ levels were revealed using anti-SepF (α-SepF) and anti-FtsZ (α-FtsZ) antibodies. **b.** Growth curves of WT cells in 1% *myo*-inositol (green triangles), *P*_*ino*_-*sepF* in the absence (blue circles) or presence (red squares) of 1% *myo*-inositol. **c.** Representative images in phase contrast (upper row) and HADA fluorescent signal (lower row) of *P*_*ino*_-*sepF* and WT strains in 1% *myo*-inositol at time points 0, 3, 6 and 9 hours after *myo*-inositol addition. Heat maps representing the localization pattern of HADA at 0, 3 and 6 hours are shown in Supplementary Fig. 2a; **d.** Violin plot showing the distribution of cell length at time points 0, 3, 6 hours after *myo*-inositol addition for *P*_*ino*_-*sepF* (top) and WT (bottom) from panel (c). The number of cells used in the analyses (n) is indicated below each violin representation, triplicate analyses are shown in Supplementary Fig. 2b. **e.** Representative images in phase contrast (upper row), HADA fluorescent signal (middle row) and mNeon-FtsZ fluorescent signal (bottom row) at time points 0, 3, 6 and 9 hours after *myo*-inositol addition. **f.** Heat maps representing the localization pattern of HADA and mNeon-FtsZ at 0, 3 and 6 hours. n numbers represent the number of cells used in the analyses. Triplicate analyses for the distribution of cell length at time points 0, 3, 6 hours as well as heatmaps for fluorescence distribution are shown in Supplementary Fig. 5. Scale Bars are 5 μm.

Using the fluorescent D-ala-D-ala analogue (HADA ^29^) to label newly incorporated peptidoglycan (PG), we showed that SepF depletion did not affect polar elongation (Fig. 1c and Supplementary Fig. 2). However, PG incorporation at mid-cell was lost and the cells were unable to form septa, thus showing that SepF is an essential component of the divisome in *Corynebacteria*. This absence of septa is clearly different from the SepF depletion phenotype in *B. subtilis*, where septa were present but largely deformed ^19^, suggesting that SepF homologues might have evolved different functions linked to the presence or absence of other auxiliary proteins such as FtsA. A phylogenetic analysis of bacterial SepF homologues shows that the proteins from *Firmicutes* and *Actinobacteria* do indeed fall into two clearly distinct groups (Supplementary Fig. 3) and suggests vertical inheritance with no horizontal transfer between both phyla. Interestingly, detectable FtsA homologues could not be identified in *Actinobacteria* nor in *Cyanobacteria* or some early-branching *Firmicutes*, which – together with the presence of SepF-like proteins (but not FtsA) in some archaeal lineages – would suggest an ancestral role for SepF in cell division.

Above we showed that septal PG synthesis was impaired in the absence of SepF, indicating that the cells could no longer assemble a functional divisome at mid-cell. As the Z-ring precedes septum formation, we asked what happened to FtsZ localization during depletion. We introduced mNeon-FtsZ as a dilute label (about 3% of total FtsZ levels) under the control of *P*_*gntK*_, a tight promoter that is repressed by sucrose and induced by gluconate 30 (Supplementary Fig. 4a,b). The minimal leakage of this promoter in sucrose was sufficient to give a signal for mNeon-FtsZ without affecting the phenotype, which remained wild-type-like (Supplementary Fig. 4c) and the phenotype during SepF depletion was comparable to that of the *P*_*ino*_-*sepF* strain (Supplementary Fig. 5). mNeon-FtsZ was followed every 3 hours during the depletion of SepF. As expected, mNeon-FtsZ localized to mid-cell at time point 0, when SepF was still present, in a typical “Z-ring” (Fig. 1e and Supplementary Fig. 5). From the following time point at 3 hours, mNeon-FtsZ was completely delocalized into foci all over the cell, showing that SepF is indeed necessary for bringing FtsZ to the mid-cell and to form a unique and functional Z-ring. Interestingly at 6 hours the distribution of mNeon-FtsZ, although lost at mid-cell, appears to be clustered and not randomly distributed throughout the cell (Fig. 1f). This observation points to an as yet undiscovered FtsZ spatial regulation mechanism, since the well characterized nuclear occlusion and Min systems in *E. coli* and *B. subtilis* are absent in *Actinobacteria* ^16^. Together these data show that in the absence of SepF, FtsZ cannot assemble at mid-cell anymore and is delocalized in the cell in a non-random manner.

### Molecular details of SepF interactions

To understand the mechanisms by which SepF participates in early divisome assembly we set out to characterize the molecular details of the interaction of SepF with both the membrane and FtsZ. The molecular organization of SepF is highly conserved and, like FtsA or ZipA, the protein contains an intrinsically disordered linker (L) of about 50 residues that separates the putative membrane-binding peptide (M) at the N-terminus from the FtsZ-binding core domain (C) at the C-terminus ^11,20^ (Fig. 2a). We proved that the predicted amphipathic helix at the N-terminus of *C. glutamicum* SepF did interact with lipid membranes (Supplementary Fig. 6a-c). Using tryptophan fluorescence titration, the peptide corresponding to the first 14 amino acids of SepF (SepF_M_) was shown to bind small unilamellar vesicles (SUVs) with a *Kd* of 32 (+/− 2) μM. In far-UV circular dichroism the SepF_M_ peptide in solution behaved as a random coil and only folded into an α-helix upon interaction with SUVs, a behavior similar to that seen for *B. subtilis* SepF ^11^.

**Figure 2:**
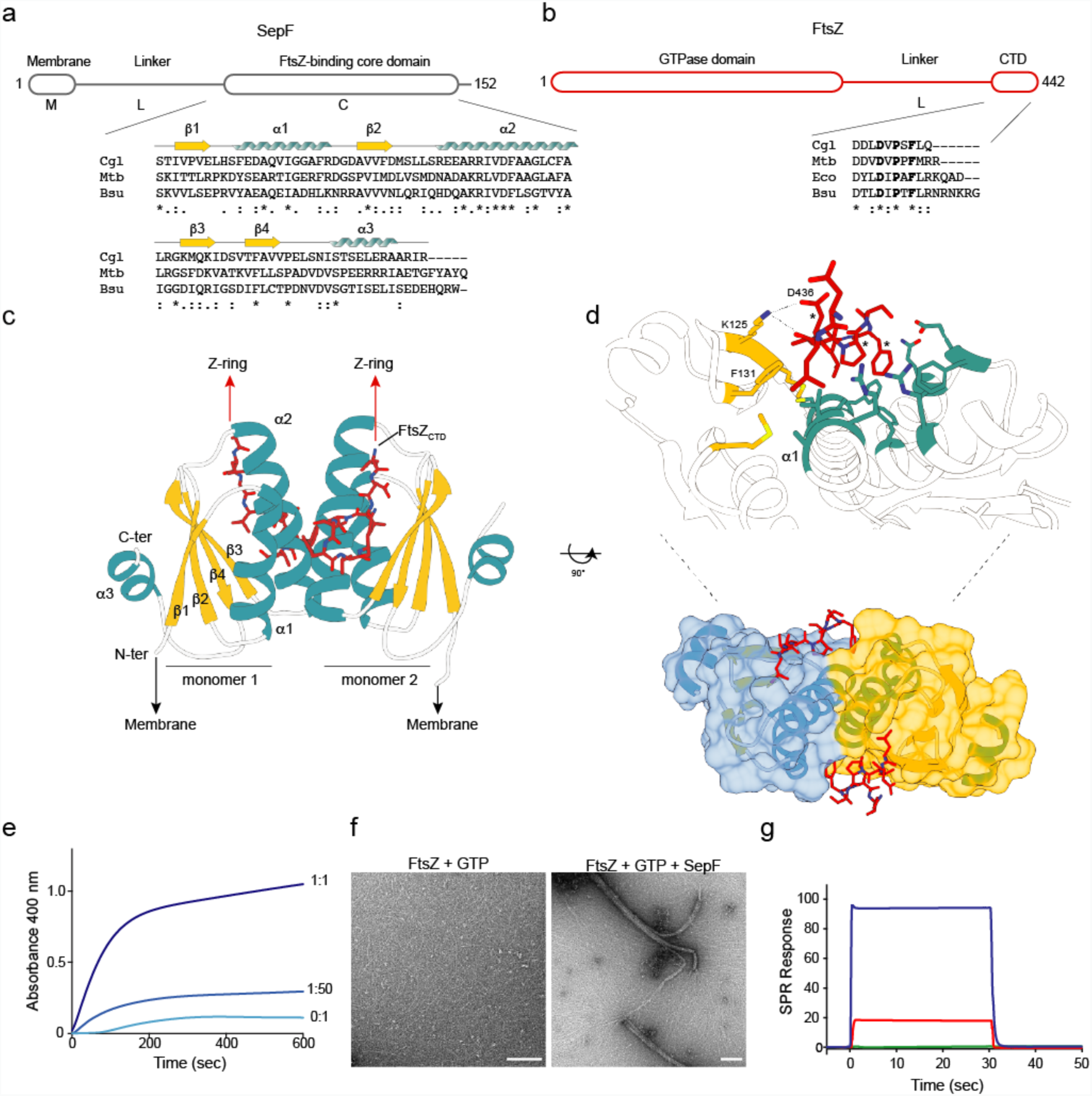
Molecular characterization of the SepF-FtsZ interaction. **a.** Schematic outline of SepF domains and sequence alignment of selected SepF homologues (Cgl, *C. glutamicum*; Mtb, *M. tuberculosis*; Bsu, *B. subtilis*). Secondary structure elements are shown above the sequences. **b.** Schematic outline of FtsZ domains and FtsZ_CTD_ sequence alignment of selected homologues (Eco, *E. coli*). Asterisks (*) indicate strictly conserved positions in the alignment and residues involved in SepF binding are shown in bold. **c.** Crystal structure of the SepF dimer in complex with two FtsZ_CTD_ peptides. The orientations of the N- and C-termini of SepF and FtsZ are compatible with membrane binding on one hand and Z-ring formation on the other. **d.** Detailed view of the FtsZ-binding pocket in SepF showing residues involved in protein-protein interactions (see Supplementary Fig. 8 for details). Conserved FtsZ_CTD_ residues D_436_, P_438_ and F_440_ are labelled (*) and SepF residues K125 and F131 were those mutated to abolish FtsZ binding. **e.** FtsZ polymerization in the presence of varying levels of SepF. The stoichiometric SepF:FtsZ ratios are indicated for each curve. **f.** Negatively stained EM micrographs of FtsZ filaments in the absence (left) and presence (right) of SepF. Scale bars are 150 nm **g.** SPR responses in resonance units (RU) for 200 μM FtsZ_CTD_ interacting with immobilized SepF (blue), SepF_F131A_ (red) and SepF_K125E/F131A_ (green).

To elucidate the structural basis of FtsZ recognition, we crystallized the SepF C-terminal core domain (SepF_ΔML_) in complex with a 10-residues peptide comprising the conserved C-terminal domain (CTD) of FtsZ (FtsZ_CTD_, Fig. 2b) and determined the crystal structure at 1.6 Å resolution (Supplementary Table 1). The SepF structure revealed a symmetric homodimer, with monomers that contain a central four-stranded β-sheet stacked against two α-helices (α1, α2) involved in dimerization and capped by a C-terminal22 α-helix (α3) on the opposite side of the sheet (Fig. 2c). The homodimer contains two identical FtsZ_CTD_-binding pockets, each made up of residues coming from both protomers (Fig. 2d), defining a dimeric functional unit for the SepF-FtsZ interaction. This 2:2 binding stoichiometry can explain mechanistically why *B. subtilis* SepF has a bundling effect on FtsZ protofilaments ^31^. Corynebacterial SepF has a similar capability, as showed by FtsZ polymerization assays at different SepF concentrations (Fig. 2e). Even at sub-stoichiometric amounts of full-length SepF, the data showed an immediate influence on polymerization dynamics and a strong FtsZ bundling effect. Comparable changes on FtsZ polymerization were also observed for the C-terminal core alone (SepF_ΔML_) but not for a SepF double mutant (SepF_K125E/F131A_, see below) that is unable to bind FtsZ (Supplementary Fig. 7), excluding the possibility that the light scattering signal could result from SepF polymerization alone. Furthermore, visualization of the protein mixture by negative stain electron microscopy (EM) after 10 minutes of incubation clearly showed thick bundles of FtsZ protofilaments as well as highly curved filaments (Fig. 2f).

The bound FtsZ_CTD_ is clearly visible in the electron density map (Supplementary Fig. 8a) and adopts a hook-like extended conformation that fits into a mostly hydrophobic binding pocket formed by conserved SepF residues, where it is further stabilized by additional intermolecular hydrogen-bonding interactions (Fig. 2d and Supplementary Fig. 8b). The apparent *Kd* value for the SepF_ΔML_-FtsZ_CTD_ interaction, as determined by surface plasmon resonance (SPR), was 15 μM (+/− 1 μM) (Supplementary Fig. 9), a value that is in the same range as those previously reported for other FtsZ_CTD_ interactors such as ZipA^32^, FtsA^33^ or ZapD^34^. The interface was further validated by mutating two FtsZ_CTD_-contact residues in SepF: F131A and K125E. Compared with the wild-type protein, the single mutant SepF_ΔML,F131A_ was greatly compromised for FtsZ binding (apparent *Kd* = 340 +/− 47 μM), whereas the double mutant SepF_ΔML,K125E/F131A_ exhibited no detectable binding in the range of protein-peptide concentrations tested (Fig. 2g and Supplementary Fig. 9).

Most FtsZ-binding proteins that have been characterized to date recognize the FtsZ_CTD_, which represents a ‘landing pad’ for FtsZ interactors ^35^. Other known structures of regulatory proteins in complex with FtsZ_CTD_ include *T. maritima* FtsA and the *E. coli* proteins ZipA, SlmA and ZapD 32-34,36. These crystal structures had shown that the FtsZ_CTD_ peptide can adopt multiple conformations depending on its binding partner, from full- or partial-helical states as in the FtsA or ZipA complexes to distinct extended conformations as in ZapD or SlmA. The SepF-bound structure of the FtsZ_CTD_ peptide revealed yet another non-helical conformation, reflecting the large conformational space that this small, highly conserved sequence can adopt in different biological contexts. It is interesting to note that SlmA and SepF, despite their different structures and binding pockets, interact with the same highly conserved hydrophobic motif of the FtsZ_CTD_ (Supplementary Fig. 10).

### Membrane and FtsZ binding *in vivo*

To further evaluate the physiological roles of SepF-membrane and SepF-FtsZ interactions we constructed fluorescently tagged SepF constructs that were either unable to bind the lipid membrane (SepF_ΔML_-Scarlet) or impaired for FtsZ binding (SepF_K125E/F131A_-Scarlet). When SepF-Scarlet was overexpressed under the control of the *P*_*gntK*_ promoter, it partially complemented the SepF depletion strain for growth even though the cells were more elongated than control cells (Figs. 3a,b and Supplementary Fig. 11). In contrast, both SepF_ΔML_-Scarlet and SepF_K125E/F131A_-Scarlet completely failed to complement the strains and showed distinct localization patterns (Fig. 3c). Importantly, the SepF_K125E/F131A_-Scarlet construct containing the mutations that abolish FtsZ interaction was diffuse in the cytoplasm, showing that FtsZ binding is needed for membrane attachment even though the membrane-binding sequence is present in the SepF construct. Previous work had shown that SepF from *M. smegmatis* was dependent on FtsZ for localization at the Z-ring ^20^, suggesting that a dynamic interplay between the two proteins is required for membrane association and Z-ring formation, and that SepF needs to act in the early actinobacterial divisome. The same behavior was reported for FtsA from *E. coli*, which also requires FtsZ for localization at the septum ^37^.

**Figure 3:**
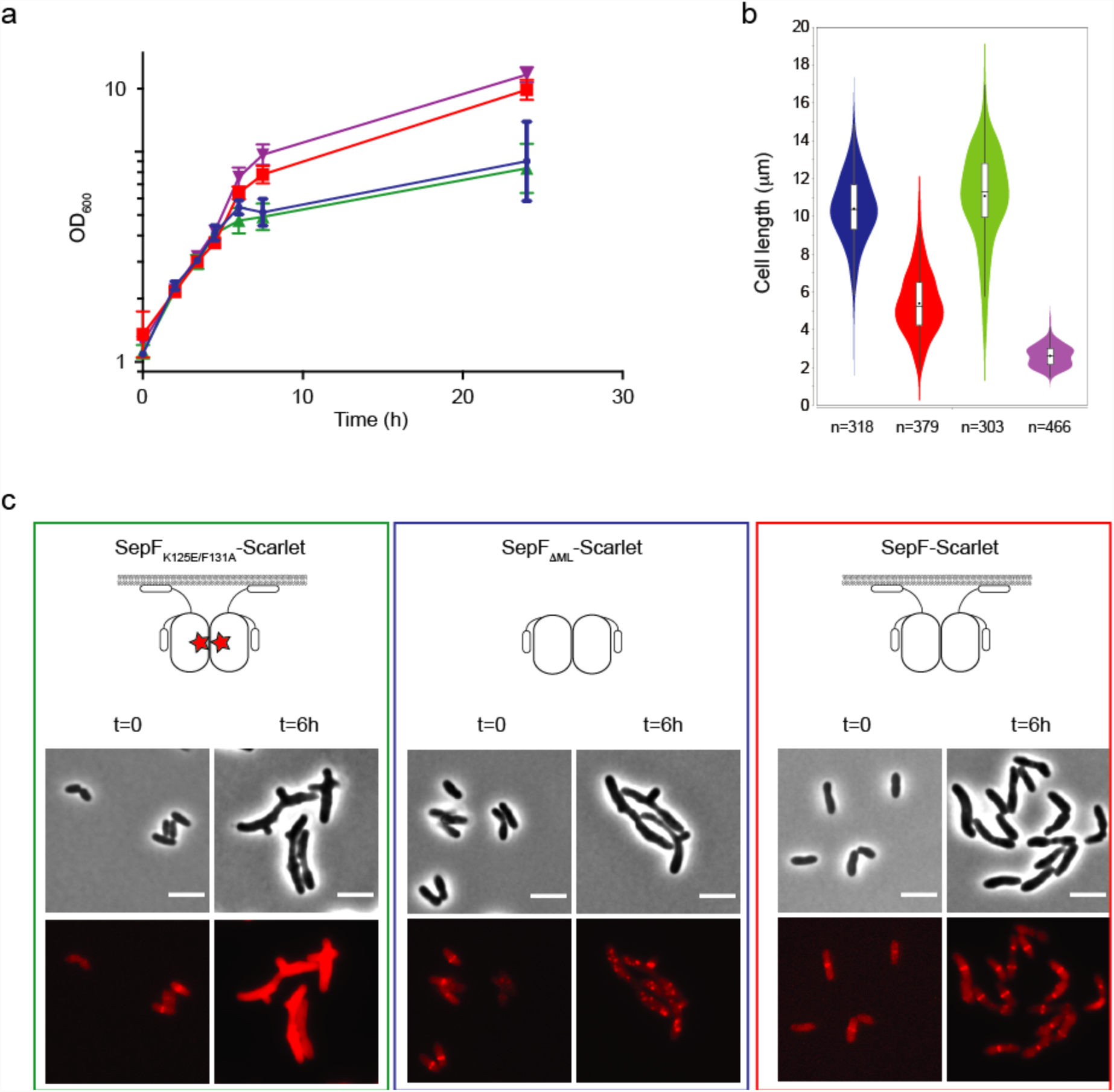
Complementation and localization of SepF mutants in the *P*_*ino*_-*sepF* strain. **a.** Growth curves of SepF-Scarlet (red), SepF_ΔML_-Scarlet (blue), SepF_K125E/F131A_-Scarlet (green) *expressed in the P*_*ino*_-*sepF-P*_*gntK*_ background in 1% *myo*-inositol plus 1% gluconate and *P*_*ino*_-*sepF-P*_*gntK*_ strain in 0% *myo*-inositol plus 1% gluconate (purple). **b.** Violin plot showing the distribution of cell length at time point 6 hours after *myo*-inositol and gluconate addition for the strains corresponding to the growth curve (same colour code). **c.** Representative images in phase contrast (upper row) and Scarlet fluorescent signal (lower row) of the complemented strains of (**a**). t=0 corresponds to the strains before depletion by *myo*-inositol and induction of exogenous *P*_*gntK*_ controlled constructs by gluconate. Western blots of whole cell extracts from the above strains during depletion as well as triplicate analyses for the distribution of cell length at time points 6 hours are shown in Supplementary Fig. 11. Scale bars = 5 μm

When we removed the membrane-binding and linker domains in the SepF_ΔML_-Scarlet construct, we observed distinct foci of SepF, similar to those seen for mNeon-FtsZ. The SepF_ΔML_-Scarlet construct can still bind and bundle FtsZ as shown above but has lost the capacity to find the mid-cell and go to the membrane (Fig. 3c). Together the above results demonstrate that SepF and FtsZ are intimately linked and interdependent to form a functional Z-ring and a viable cell in *Actinobacteria*.

### A putative regulatory role for the C-terminal helix of SepF

The crystal structure of the globular core of SepF reported in this work differs from the available structures of other bacterial and archaeal SepF-like homologues in the PDB (codes 3P04, 3ZIE, 3ZIG, 3ZIH) in that it contains an additional helix (α3) at its C-terminus (Fig. 2c). This helix was predicted but not seen in the other structures because it is either missing in the construct or structurally disordered in the crystal ^11^. Helix α3 formation and stabilization were not due to FtsZ binding, because the crystal structure of unliganded SepF_ΔML_ (Supplementary Table 1) revealed the same overall structure than FtsZ_CTD_-bound SepF_ΔML_ (rmsd of 0.83 Å for 160 equivalent Cα atoms in the homodimer). The presence of this helix has important functional implications as it caps the β-sheet (Fig. 4a) that was previously described as the dimerization interface of *B. subtilis* SepF ^11^. In fact, the crystal structure of the latter (and those of SepF-like proteins from *Pyrococcus furiosus* and *Archaeoglobus fulgidus*) revealed two possible dimerization interfaces: one that occurred *via* the central β-sheet, which was proposed to define the functional unit, and one that occurred *via* a 4-helix bundle, thought to be involved in protein oligomerization and ring formation ^11^ (Supplementary Fig. 12a). Our crystal structure superimposed well with the 4-helix bundle-mediated SepF dimers from *B. subtilis*, *P. furiosus* and *A. fulgidus*, confirming that the *C. glutamicum* SepF homodimer (Fig. 2c) is conserved across different bacterial and archaeal species.

As the functional SepF unit is conserved in *C. glutamicum* and *B. subtilis*, the β-sheet-mediated interface observed in the other SepF crystals could therefore mediate protein polymerization in solution. This interaction is precluded in the *C. glutamicum* SepF structure by the presence of the C-terminal helix α3 (Fig. 4a). Interestingly, inspection of the unliganded SepF_ΔML_ structure revealed that this helix displays considerably higher B-factor values than the rest of the protein (Fig. 4b) and a similar trend is also observed in the SepF_ΔML_-FtsZ_CTD_ complex, suggesting that helix α3 could play a regulatory role on SepF polymerization by uncovering the outer face of the β-sheet for intermolecular interactions. To further investigate this hypothesis, we removed the helix and crystallized the resulting SepF_ΔML,Δα3_ construct alone and in complex with FtsZ_CTD_ (Supplementary Table 1). Despite a different crystal packing, the two structures did show a dimer-dimer association mediated by the opposing β-sheets in the crystal, generating linear SepF polymers similar to those observed for the *B. subtilis* homologue (Fig. 4c and Supplementary Fig. 12). These observations demonstrate that the β-sheet is indeed prone to self-interaction in solution and suggest that the amphipathic helix α3 might regulate these interactions, possibly by engaging itself in other intermolecular interactions. It is interesting to note, however, that the anti-parallel orientation of the interacting β-sheets in the three crystal forms of *C. glutamicum* SepF, although identical to each other, differs from the parallel orientation seen in the *B. subtilis* SepF structure, thus pointing to species-specific polymerization mechanisms.

**Figure 4:**
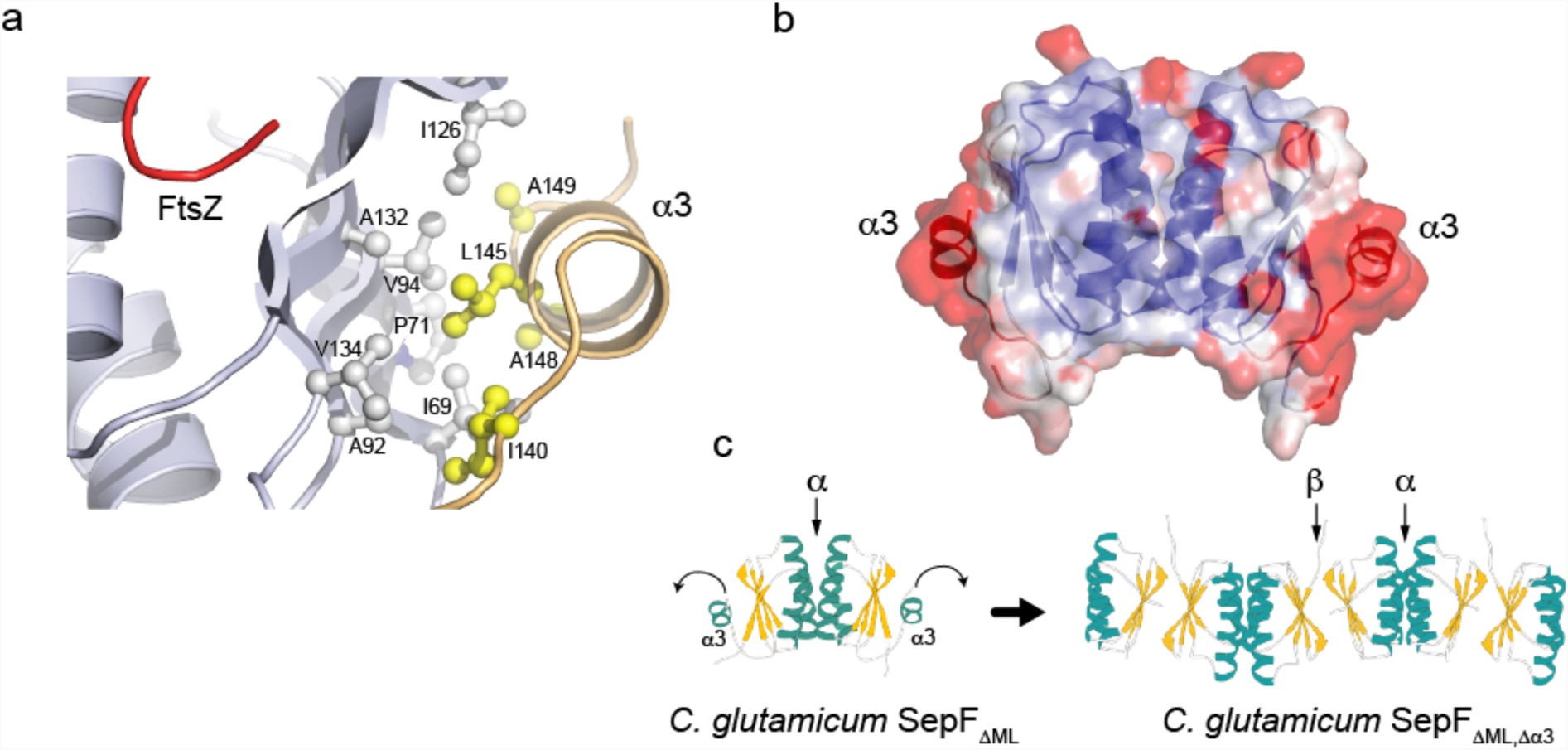
Putative regulatory role of helix α3. **a.** Hydrophobic interactions mediate the association of the C-terminal helix α3 (yellow) with the central β-sheet (grey) in the SepF protomer. **b.** Overall structure of the unliganded SepF_ΔML_ homodimer color coded according to temperature factors, from blue (lowest values) to red (highest values). **c.** Deletion of helix α3 in SepF_ΔML, Δα3_ promotes the tight interaction between opposite β-sheets from different SepF dimers, leading to the formation of linear SepF polymers (see also Supplementary Fig. 12).

The above polymerization model can account for the formation of SepF rings in *B. subtilis* ^22^. However, *C. glutamicum* SepF devoid of helix α3 (i.e. with the β-sheet accessible for protein-protein interactions) remained dimeric in solution and no higher order assemblies could be detected by size exclusion chromatography or analytical ultracentrifugation (Supplementary Fig. 13). Moreover, while we could observe regular rings of ~40 nm diameter in negative stain EM for the related SepF homologue from *M. tuberculosis* (Supplementary Fig. 14), our extensive attempts at detecting ring-like structures of *C. glutamicum* SepF were unsuccessful. In this particular case, oligomerization may therefore require an increased local concentration, as found for instance in crystallization conditions or in a physiological situation linked to lipid membrane and/or FtsZ filament interactions, where avidity effects may push the equilibrium towards the oligomeric state. Taken together the above biophysical data suggest that helix α3 could regulate the oligomeric state of SepF by controlling β-sheet-mediated interactions during Z-ring formation and divisome assembly.

### SepF oligomerization occurs upon membrane binding and is reversed by FtsZ interactions

We next investigated the effects of lipid membranes and FtsZ-binding on SepF polymerization. We found that incubation of SepF with SUVs led to SepF concentration-dependent polymerization in turbidity assays (Fig. 5a and Supplementary Fig. 15) and to the formation of large structures in dynamic light scattering (DLS) experiments (Supplementary Figs. 16a-c). When we looked at the end point of these reactions in negative stain EM the lipid vesicles were tubulated (Fig. 5a), indicating that the N-terminal amphipathic helix of SepF has the capability to induce membrane remodeling upon protein polymerization. Surprisingly this behavior was completely reversed when the FtsZ_CTD_ peptide was added to the reaction (Fig. 5b and Supplementary Fig. 16d): SepF depolymerized, lipid vesicles recovered their initial size in DLS, and small regular vesicles with no tubulation were observed in electron micrographs. When the same experiment was carried out with SepF_Δα3_, lacking the regulatory C-terminal helix, polymerization and vesicle tubulation were also seen but could only be partially reversed after addition of the FtsZ_CTD_ peptide (Supplementary Fig. 17).

**Figure 5:**
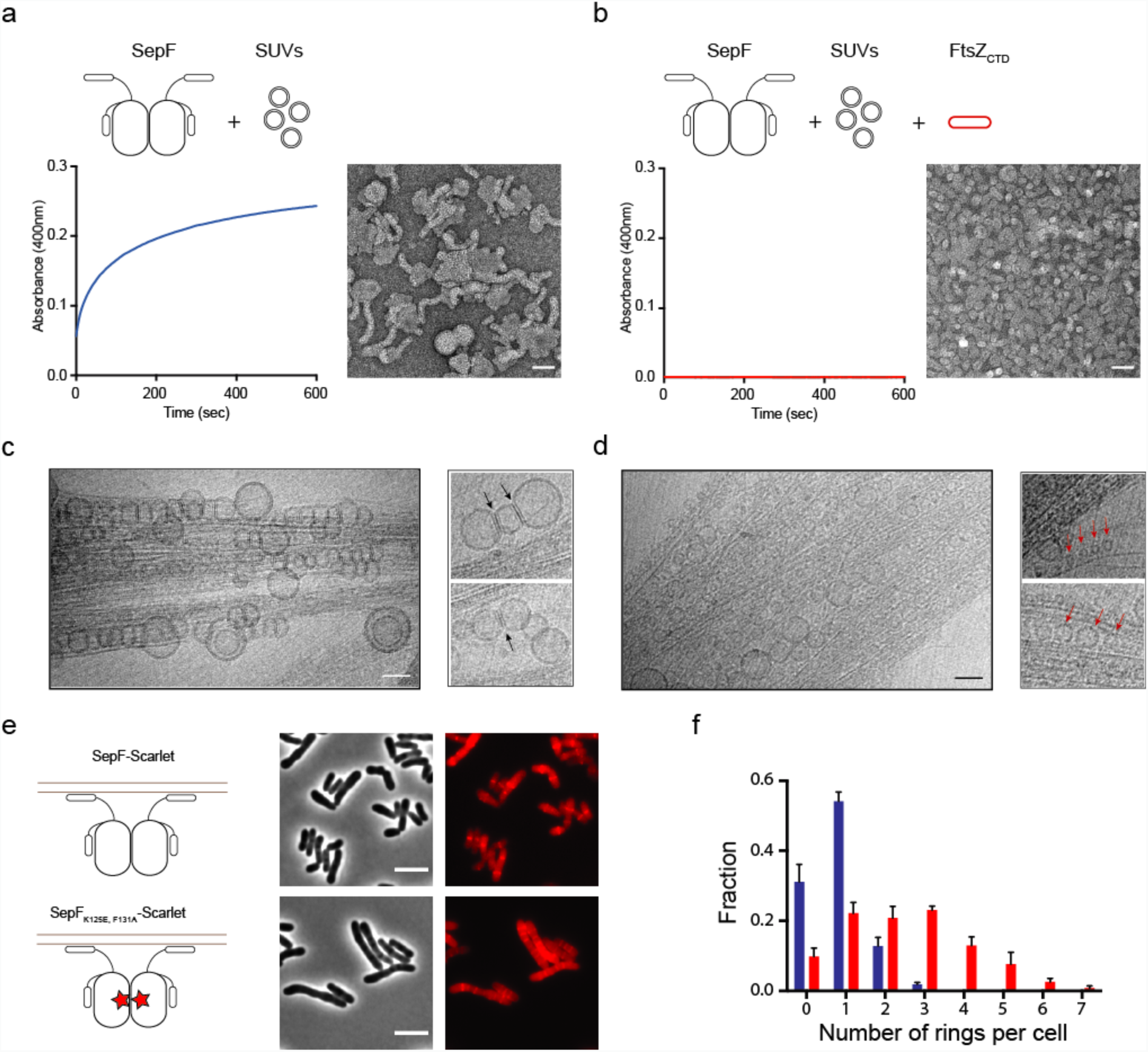
SepF oligomerization upon lipid membrane-binding and reversal by FtsZ. **a.** Polymerization assay and negatively-stained EM images of the end-point of the assay for SepF in the presence of SUVs. **b.** Idem for SepF in the presence of SUVs + FtsZ_CTD_. **c.** Cryo-EM images of the ternary complex full-length FtsZ-GTP + SepF + SUVs. A bundle of FtsZ filaments is decorated by tubulated lipid vesicles, mostly found on the periphery of the bundles where fewer FtsZ binding sites are available. The smaller panels show two examples of tubulated lipid vesicles, where black arrows indicate the putative SepF rings. Scale bar, 50 nm. **d.** In the central regions of the bundles, several small, non-tubulated, vesicles are in close contact with FtsZ filaments. Red arrows indicate a few examples of these vesicles in the right panels. **e.** Phase contrast (left columns) and Scarlet fluorescent signal (right column) images show the phenotypic differences observed upon overexpression of SepF-Scarlet (top) and SepF_K125E/F131A_-Scarlet (bottom) in the WT strain, where endogenous SepF is present. Western blots of whole cell extracts from the above strains as well as triplicate analyses for the distribution of cell length are shown in Supplementary Fig. 18. Scale bars = 5 μm. **f.** Frequency histogram indicating the number of ring-like structures per cell for SepF-Scarlet (blue) and SepF_K125A/F131A_-Scarlet (red). These analyses were carried out in triplicates of 336<n<477 cells per experiment and error bars are shown in the graph.

In cryo-EM experiments with full-length FtsZ (instead of FtsZ_CTD_), we have been able to trap an intermediate state of the ternary complex (lipid membranes, SepF, FtsZ-GTP). In the images (Fig. 5c) lipid vesicles of different sizes are brought together by rings of SepF (these membranes are negatively charged and do not form these structures by themselves). The fused large vesicles can only be found at the periphery of the FtsZ filaments, where fewer FtsZ_CTD_ tails are available for binding. On the other hand, for vesicles trapped in the central bundle regions the tubular structures are lost. In an end-point scenario (Fig. 5d) all the lipid vesicles are small and decorate the FtsZ filaments, showing that SepF still binds to lipids but cannot polymerize (and tubulate) anymore. Taken together, these results point to a dynamic link between SepF oligomerization and membrane remodeling coupled to an antagonistic effect upon FtsZ binding, where the SepF C-terminal helix (α3) would act as a regulatory switch.

Further evidence for the interdependence of SepF and FtsZ dynamics *in vivo* was obtained by overexpressing SepF_K125E/F131A_-Scarlet (which is unable to bind FtsZ) in the WT background. From mass spectrometry data (immunoprecipitation experiments coupled to crosslinking) we found that FtsZ binding in the strain expressing SepF_K125E/F131A_-Scarlet was greatly reduced (7.2 fold; see Methods section and Supplementary Table 2) compared to SepF-Scarlet, but not abolished, showing that the mutant and endogenous SepF proteins interacted with each other. When we looked at the localization of SepF in these two strains, we observed a net increase in the number of ring-like structures per cell for SepF_K125E/F131A_-Scarlet as opposed to mostly single rings at mid-cell for SepF-Scarlet (Figs. 5e,f and Supplementary Fig. 18). Thus, a corrupted FtsZ-binding site led to the formation of multiple, more stable SepF rings that contain not only fully functional endogenous SepF but also non-functional SepF_K125E/F131A_-Scarlet. Since recombinant SepF levels exceed the endogenous protein levels (Supplementary Fig. 18), these partially functional rings are expected to contain patches that cannot bind and stabilize FtsZ filaments. This would in turn interfere with the formation of a fully dynamic oligomeric Z-ring structure, which requires correct alignment and stabilization for solid treadmilling-driven assembly of the division machinery ^10,38,39^. The fact that dynamics between FtsZ and SepF are affected in these mutants would mean that the rings of SepF become more stable and dissociate less from the membrane.

FtsZ by itself has the properties not only to self-organise and provide directionality, but also to deform lipids when a membrane binding motif is attached to the protein ^40–42^. In this case, FtsZ has been shown to assemble into dynamic vortices *in vitro* without the need for accessory proteins such as FtsA, but critically relying on concentration thresholds ^41^. Concentration dependence is also true for accessory proteins such as the bundling protein ZipA ^43^, or ZapA ^44^ where the effect on FtsZ dynamics heavily relies on the protein concentrations used. In the bacterial cell FtsZ does not contain a membrane binding domain and the self-assembly is counteracted by several parameters such as molecular crowding ^45,46^ as well as spatial and temporary constraints for mid-cell localization. A functional tethering system thus needs to allow for accumulation of enough FtsZ molecules (i.e. *via* bundling) without trapping these bundles in static states. In *E. coli*, it was recently shown that the bundling protein ZapA increases FtsZ filament stability without affecting the treadmilling activity ^44^. The Z-ring also needs to constantly adapt to the dynamics and shrinking of the membrane invagination during septal closure, and dynamic exchange is thus primordial to all FtsZ interacting systems.

Our results point to a dual role of corynebacterial SepF during the early assembly of the divisome (Fig. 6). In its dimeric form SepF fulfills FtsZ bundling and membrane tethering functions. On the other hand, SepF polymerization has a membrane remodeling effect, antagonized by FtsZ binding, which might contribute to septum formation during cell division. A dynamic interplay between these two roles would thus be required for Z-ring assembly. Membrane remodeling activity has recently been described for the *E. coli* membrane tether FtsA ^25^. For SepF a plausible scenario could be that, during the early stages of assembly, polymerized FtsZ fragments are bundled and tethered to the membrane mainly by dimeric SepF, which – possibly assisted by auxiliary regulatory factors yet to be identified – will help FtsZ polymers to stabilize, find directionality, and start treadmilling to form a functional Z-ring. At the same time, this process would increase the local concentration of SepF at the membrane. Both treadmilling and bundling contribute to remove available FtsZ_CTD_ binding sites, leading to SepF polymerization and membrane invagination, contributing to the net force required for cell constriction ^47^. A possible consequence of this model is that SepF-induced septum formation would only occur when enough FtsZ has been accumulated at the membrane and treadmilling starts, making of SepF a checkpoint protein that would initiate constriction only once the cytomotive machinery is fully functional.

**Figure 6:**
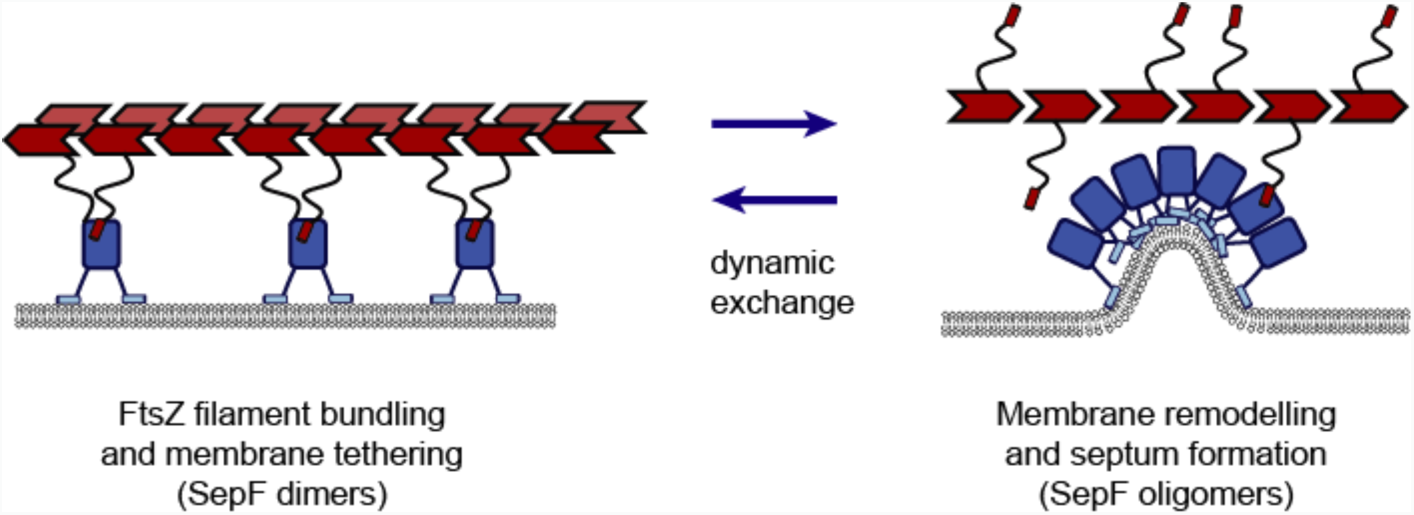
Schematic model depicting the possible role of dynamic SepF oligomerization during Z-ring assembly and septum formation. Dimeric SepF (blue) can bind and bundle FtsZ protofilaments (red) and tether them to the membrane (left panel). An increasing concentration of both SepF and FtsZ will help FtsZ polymers to stabilize, find directionality, and start treadmilling to form a functional Z-ring. Concomitantly, this process will increase the local concentration of SepF at the membrane. Both treadmilling and bundling will contribute to remove available FtsZ_CTD_ binding sites, prompting SepF to polymerize and remodel the membrane contributing to membrane invagination, required during the early stages of septum formation (right panel).

## Methods

### Bacterial strains and growth conditions

All bacterial strains used in this study are listed in the Supplementary Table 3. *Escherichia coli* DH5α or CopyCutter EPI400 were used for cloning and were grown in Luria-Bertani (LB) broth or agar plates at 37°C supplemented with 50 µg/ml kanamycin when required. For protein production, *E. coli* BL21 (DE3) was grown in LB or 2YT broth supplemented with 50 µg/ml kanamycin at the appropriate temperature for protein expression.

#### Corynebacterium glutamicum

ATCC 13032 was used as a wild-type (WT) strain. *C. glutamicum* was grown in brain heart infusion (BHI) at 30°C and 120rpm and was supplemented with 25 µg/ml kanamycin when required. When specified, minimal medium CGXII ^48^ supplemented with sucrose, gluconate, tetracycline, kanamycine and/or *myo*-inositol was used.

### Cloning and mutagenesis for recombinant protein production in *E. coli*

The genes encoding for *C. glutamicum sepF* (*cg2363*), *M. tuberculosis sepF (Rv2147c*), and *C. glutamicum ftsZ (cg2366*) were codon optimized and synthesized for *E. coli* protein production (Genscript) and used as templates for subsequent cloning. They were cloned into a pT7 vector containing an N-terminal 6xHis-SUMO tag. SepF mutants containing different domains: SepF_ΔML_ (amino acids 63 to 152), SepF_ΔML,Δα3_ (amino acids 63 to 137), SepF_Δα3_ (amino acids 1 to 137), SepF_ΔML,F131A_, SepF_ΔML,K125E/F131A_, SepF_K125E/F131A_ and *M. tuberculosis* SepF_ΔML_ (*Mtb*SepF_ΔML_, comprising amino acids 122 to 218) were constructed using either pairs of complementary primers carrying the desired deletion or point mutations on the primers listed in the Supplementary Table 4. PCR products were digested with DpnI and transformed into chimio-competent *E. coli* cells. All plasmids were verified by Sanger sequencing (Eurofins Genomics, France).

### Cloning and mutagenesis for recombinant protein expression in *C. glutamicum*

For ectopic recombinant expression of the different constructs in *C. glutamicum*, we modified the synthetic pTGR5 shuttle expression vector 49 by exchanging the *P*_*tac*_ promoter present in the plasmid by the promoters *P*_*gntK*_ of *C. glutamicum* or *P*_*tet*_ from pCLTON1 vector 50 yielding plasmids pUMS_3 and pUMS_40 respectively (Supplementary Table 3). FtsZ and SepF variants were assembled in these plasmids by either Gibson assembly or site-directed mutagenesis using the primers listed in Supplementary Table 4. For cellular localization studies, SepF and FtsZ were fused to mScarlet-I ^51^ or mNeonGreen ^52^ respectively, including a flexible linker between the fluorescent protein and the gene of interest.

### Construction of the *sepF* conditional mutant

To obtain the conditional depletion of *sepF*, the endogenous *sepF* was placed under the control of a repressible promoter. Using the two-step recombination strategy with the pk19mobsacB plasmid, we inserted the native promoter of the inositol phosphate synthase gene (*ino1* – *cg3323*) which can be repressed in the presence of *myo*-inositol ^27^ A terminator followed by the *ino1* promoter was amplified by PCR from pk19-P3323-*lcpA* ^53^. The 500bp up-stream region and down-stream region of *sepf* were amplified using chromosomal DNA of *C. glutamicum* ATCC 13032 as a template. The different fragments were assembled in a pk19mobsacB backbone by Gibson assembly (NEB). The plasmid was sequenced and electroporated into WT *C. glutamicum* ATCC13032. Positive colonies were grown in BHI media supplemented with 25 µg/ml kanamycin over night at 30°C and 120 rpm shaking. The second round of recombination was selected by growth in minimal medium CGXII plates containing 10% (w/v) sucrose. The insertion of the *ino1* promoter was confirmed by colony PCR and sequenced (Eurofins, France). All the oligonucleotides used in order to obtain and check this strain are listed in the Supplementary Table 4.

### Growth curves

All strains were plated in CGXII media plates with 4% (w/v) sucrose as a carbon source for 2 days at 30°C and then single colonies were inoculated in 10 ml of CGXII media 4% (w/v) sucrose overnight at 30°C and 120 rpm shaking. The next day, 20 ml of CGXII media supplemented with the appropriate repressor or inducer were inoculated with the overnight cultures to a starting OD_600_ of 1. OD_600_ measurements were taken every 1.5 hours. Each growth curve represents the average of 3 different growth curves originally from 3 different single colonies. For each time point a sample for western blot was taken. When required, CGXII media 4% (w/v) sucrose was supplemented with either 1% (w/v) *myo*-inositol, 50 ng/ml tetracycline or 1% (w/v) gluconate and 25 μg/ml kanamycin.

### Protein expression and purification

N-terminal 6xHis-SUMO-tagged SepF, and derivate mutants (both from *C. glutamicum* and *M. tuberculosis*) were expressed in *E. coli* BL21 (DE3) following an auto-induction protocol ^54^. After 4 hours at 37°C cells were grown for 20 hours at 20°C in 2YT complemented autoinduction medium containing 50 µg/ml kanamycin. Cells were harvested and flash frozen in liquid nitrogen. Cell pellets were resuspended in 50 ml lysis buffer (50 mM Hepes pH8, 300 mM NaCl, 5% glycerol, 1 mM MgCl_2_, benzonase, lysozyme, 0.25 mM TCEP, EDTA-free protease inhibitor cocktails (ROCHE) at 4°C and lysed by sonication. The lysate was centrifuged for 30min at 30.000 x g at 4°C. The cleared lysate was loaded onto a Ni-NTA affinity chromatography column (HisTrap FF crude, GE Healthcare). His-tagged proteins were eluted with a linear gradient of buffer B (50 mM Hepes pH8, 300 mM NaCl, 5% glycerol, 1 M imidazole). The eluted fractions containing the protein of interest were pooled and either dialysed directly (for SPR experiments) or dialysed in the presence of the SUMO protease (ratio used, 1:100). Dialysis was carried out at 4°C overnight in 50 mM Hepes pH8, 150 mM NaCl, 5% glycerol, 0.25 mM TCEP. Cleaved his-tags and his-tagged SUMO protease were removed with Ni-NTA agarose resin. The cleaved protein was concentrated and loaded onto a Superdex 75 16/60 size exclusion (SEC) column (GE Healthcare) pre-equilibrated at 4°C in 50 mM Hepes pH8, 150 mM NaCl, 5% glycerol. The peak corresponding to the protein was concentrated, flash frozen in small aliquots in liquid nitrogen and stored at −80°C.

Codon optimized N-terminal 6xHis-SUMO-tagged *C. glutamicum* FtsZ was produced and purified as described above, except that KCl was used instead of NaCl and a TALON FF crude column (GE Healthcare) was used for affinity chromatography. All purified proteins used in this work have been run on an SDS-PAGE and are represented in Supplementary Fig. 19.

### Crystallization

Crystallization screens were performed for the different SepF constructs and SepF-FtsZ_CTD_ complexes using the sitting-drop vapour diffusion method and a Mosquito nanolitre-dispensing crystallization robot at 18 °C (TTP Labtech, Melbourn, UK). Optimal crystals of SepF_ΔML_ (13 mg/ml) were obtained after one week in 10% PEG 800, 0.2 M Zinc acetate, 0.1 M sodium acetate. The complex SepF_ΔML_ - FtsZ_CTD_ (DDLDVPSFLQ, purchased from Genosphere) was crystallized at 10 mg/ml SepF_ΔML_ and 5.8 mg/ml of FtsZ_CTD_ (1:5 molar ratio) after 2 weeks in 100 mM sodium acetate pH 4.6, and 30% w/v PEG 4000. SepF_ΔML,Δα3_ (17 mg/ml) crystallized within 2 weeks in 0.1 M MES pH 6, 20 %w/v PEG MME 2K and 0.2 M NaCl. The SepF_ΔML,Δα3_ - FtsZ_CTD_ complex was crystallized at 17 mg/ml SepF_ΔML,Δα3_ and 9.8 mg/ml FtsZ_CTD_ (1:5 molar ratio) within 2 weeks in 0.1 M MgCl_2_, 0.1M MES pH 6.5 and 30% w/v PEG 400 buffer. Crystals were cryo-protected in mother liquor containing 33% (vol/vol) ethylene glycol or 33% (vol/vol) glycerol.

### Data collection, structure determination and refinement

X-ray diffraction data were collected at 100 K using beamlines ID30B and ID23-1 (wavelength = 0.97625 Å) at the ESRF (Grenoble, France). All datasets were processed using XDS ^55^ and AIMLESS from the CCP4 suite ^56^. The crystal structures were determined by molecular replacement methods using Phaser ^57^ and *C. glutamicum* SepF (PDB code 3p04) as the probe model. In the case of SepF_ΔML_, the structure was also independently determined by single-wavelength anomalous diffraction (SAD) phasing using Patterson methods to localize the protein-bound Zn ions (present in the crystallization buffer) with SHELXD ^58^ and automatic model building with Buccaneer from the CCP4 suite ^59^. All structures were refined through iterative cycles of manual model building with COOT ^60^ and reciprocal space refinement with BUSTER ^61^. Highly anisotropic diffraction was observed for SepF_ΔML,Δα3_ crystals, in which one of the two monomers in the asymmetric unit was largely exposed to solvent (see Supplementary Fig. 12c) and exhibited unusually high temperature factors (average B values for all main-chain atoms of the two monomers were respectively 17 Å^2^ and 58 Å^2^). The crystallographic statistics is shown in Supplementary Table 1 and representative views of the final electron density map for each structure are shown in Supplementary Fig. 20. Structural figures were generated with Chimera ^62^ or Pymol (The PyMOL Molecular Graphics System, Version 2.0 Schrödinger, LLC). Atomic coordinates and structure factors have been deposited in the protein data bank under the accession codes 6sat, 6scp, 6scq and 6scs.

### FtsZ and SepF polymerization assay

Purified FtsZ and SepF were precleared at 25.000 x g for 15min at 4°C. FtsZ with or without SepF was added to a final concentration of 15 µM each in 500 µl final volume in polymerization buffer (100 mM KCl, 10 mM MgCl_2_, 25 mM Pipes pH 6.9). The mixture was placed into a quartz cuvette with a light path of 10 mm and 2 mM GTP were added to the reaction mixture. Data acquisition started immediately using an UV-Visible Spectrophotometer (Thermo scientific Evolution 220) during 600 seconds at 25°C using 400 nm for excitation and emission and spectra with slits widths of 1 nm.

To follow the polymerization of SepF in the presence of lipids, we used SepF or SepF mutants plus SUVs at a final concentration of 50 µM each in polymerization buffer. The mixture was placed into a quartz cuvette with a light path of 10mm and data acquisition was carried out as mentioned above. When used, FtsZ_CTD_ was added at a final concentration of 100 μM.

### Small unilamellar vesicle (SUV) preparation

Reverse phase evaporation was used to prepare small unilamellar vesicles (SUVs). A 10 mM lipids chloroform solution made of an 8:2 mixture of 1-palmitoyl-2-oleoyl phosphatidylcholine (POPC) and 1-palmitoyl-2-oleoylglycero-3-phosphoglycerol (POPG) (Avanti Polar Lipids). Chloroform was removed by evaporation under vacuum conditions and the dried phospholipid film was resuspended in a mixture of diethyl ether and buffer (25 mM Hepes pH 7.4, 10 mM MgCl_2_ and 150 mM KCl). Remaining diethyl ether was eliminated by reverse phase evaporation and by slowly decreasing the pressure to the vacuum. SUVs were obtained by sonication during 30 minutes at 4°C. The diameter and charge of SUV were determined by measuring dynamic light scattering (DLS) and electrophoretic mobility profiles on a Zetasizer Nano instrument (Malvern Instruments).

### Far-UV circular dichroism

The secondary structure of SepF_M_ in the presence or absence of SUVs was determined using synchrotron radiation circular dichroism (SRCD) carried out at the beamline DISCO (SOLEIL, Gif-sur-Yvette, France). Three individual scans were averaged to obtain final far-UV spectra. Measurements were made at 25°C with an integration time of 1.2s and bandwidth of 1nm. SRCD spectra in the far-UV (190 to 250nm) were recorded using QS cells (Hellma, France) with a path length of 100 µm. SepF_M_ was used at 100 μM in CD buffer (25 mM Hepes pH 7.4, 100 mM KCl) in the presence and absence of SUVs (POPC:POPG 8:2) at a final concentration of 2 mM. As a blank, the CD buffer was used in the absence or presence of SUVs, and subtracted from far-UV CD spectra. Finally, BestSel ^63^ was used in order to estimate the content of secondary structure.

### Lipid peptide interaction (Tryptophan fluorescence emission titration)

To estimate the partition coefficient (K_x_) between SUVs and SepF, we used the synthetic peptide MSMLKKTKEFFGLAW (purchased from Genosphere), which contains the SepF_M_ sequence with an extra W residue at the C-terminal. K_x_ is defined as the ratio of peptide concentration in the lipid and in the buffer phases. K_x_ can be expressed by the following equation:

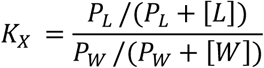

In which P_W_ represent the concentration of soluble peptide (in aqueous phase) and P_L_ the peptide concentration bound to lipid membranes (lipidic phase). [L] refers to the lipid concentration and [W] refers to the water concentration. K_x_ is directly related to the apparent dissociation constant as K_x_ * K_D_= [W] with K_D_* P_L_ = P_W_ * [L]. The equation was fitted to the experimental data using the KaleidaGraph software.

We used a FP-8200 (Jasco, Tokyo, Japan) spectrophotometer equipped with a thermostatic Peltier ETC-272T at 25°C. Experiments were performed in a cuvette 109.004F-QS (Hellma, France) containing a magnetic stirrer. A bandwidth of 5 nm was used for emission and excitation. 1 μM of peptide was used in Titration buffer (100 mM KCl and 25 mM Hepes pH 7,4). We measured the florescence emission between 300 and 400 nm at a scan rate of 125 nm/min with an excitation wavelength of 280 nm. The obtained spectra were corrected by blank subtraction (SUV light scattering in titration buffer). Next, the maximum wavelength value (λmax) and the fluorescence ratio 330/370 were calculated to measure the partition coefficient (K_x_).

### Real-time Surface Plasmon Resonance (SPR)

All experiments were carried out on a Biacore T200 instrument (GE Healthcare Life Sciences) equilibrated at 25°C in 25 mM HEPES pH 8, 150 mM KCl, 0.1 mM EDTA. For SPR surface preparation His6-Sumo tagged constructs of SepF_ΔML_, SepF_ΔML,F131A_, SepF_ΔML,K125E/F131A_ were covalently immobilized on 3 independent flow-cells of an NTA sensorchips as previously described ^64^ (GE Healthcare Life Sciences). The final immobilization densities of SepF variants ranged from 3000 to 4000 resonance units (RU; 1RU ≈ 1 pg/mm^2^).

For SPR binding assays, different concentrations of the FtsZ_CTD_ peptide (ranging from 1.56 to 400 µM) were injected sequentially at 50 µl/min on the SepF-functionalized surfaces. Association was monitored for 30 s, followed by a buffer wash for 120 s during which the full dissociation of the SepF-FtsZ_CTD_ complex was observed. The concentration dependence of the SPR steady-state signals (R_eq_) was analysed, allowing to determine the equilibrium dissociation constant K_d_, by fitting the dose/response curve with the equation R_eq_ = R_max_ * C / K_d_ + C (where C is the concentration of FtsZ_CTD_ and R_max_ the response at infinite peptide concentration).

### Phase contrast and fluorescence microscopy, Image analysis

For imaging, cultures of *C. glutamicum* were grown in BHI or minimal medium CGXII during the day and washed and inoculated into CGXII media for overnight growth. The following day cultures were diluted to OD_600_=1 and grown to the required OD (early exponential phase) for imaging. For HADA labelling, cultures were incubated with 0.5 mM HADA for 20 min at 30°C in the dark. 2% agarose pads were prepared with the corresponding growth medium and cells were visualized using a Zeiss Axio Observer Z1 microscope fitted with an Orca Flash 4 V2 sCMOS camera (Hamamatsu) and a Pln-Apo 63X/1.4 oil Ph3 objective. Images were collected using Zen Blue 2.6 (Zeiss) and analyzed using the software Fiji ^65^ and the plugin MicrobeJ ^66^ to generate violin plots and fluorescent intensity heat maps. The experiments were performed as biological triplicates.

### Western Blot

To prepare cell extracts, bacterial cell pellets were resuspended in buffer lysis (50 mM Bis-Tris pH 7,4; 75 mM 6-Aminocaproic Acid; 1 mM MgSO4; Benzonase and protease Inhibitor) and disrupted at 4 °C with 0.1 mm glass beads and using a PRECELLYS 24 homogenizer. Crude extracts (120μg) were analysed by SDS-PAGE, electrotransferred onto a 0,2μm Nitrocellulose membrane and blocked with 5% (w/v) skimmed milk. Membranes were incubated with an anti-SepF antibody (produced by Covalab, Supplementary Fig. 21a) or an anti-FtsZ antibody (produced by Proteogenix, Supplementary Fig. 21b) for 1 h at room temperature. After washing in TBS-Tween buffer (Tris-HCl pH8 10 mM; NaCl 150 mM; Tween20 0,05% (vol/vol), the membrane was incubated with an anti-rabbit horseradish peroxidase-linked antiserum (GE healthcare) for 45 minutes. The membrane was washed and revealed with HRP substrate (Immobilon Forte, Millipore) and imaged using the ChemiDoc MP Imaging System (BIORAD).

### Electron microscopy

For negative stain sample preparations, incubations were performed at room temperature. SUVs and SepF constructs (50 μM) were incubated in polymerization buffer (100 mM KCl, 10 mM MgCl_2_, 25 mM Pipes pH 6.9) for 10 minutes. When used, FtsZ_CTD_ was added at a final concentration of 100 μM. To image the FtsZ polymers with and without SepF, we used a final concentration of protein at 21 μM and 13 μM respectively in EM buffer containing 50 mM HEPES pH 7.4, 300 mM KCl and 10 mM MgCl_2_ supplemented with 3 mM GTP. FtsZ was incubated with or without SepF at room temperature and imaged after 10 minutes.

To image SepF rings form *M. tuberculosis*, *Mtb*SepF_ΔML_ was used at a final concentration of 50 μM in buffer containing 50 mM HEPES pH 7.4, 300 mM KCl and 10 mM MgCl_2_. For all samples, 400 mesh carbon coated grids (Electron Microscopy Sciences; CF 400-Cu) were glow-discharged on an ELMO system for 30 sec at 2mA. 5 μl samples were applied onto the grids and incubated for 30s, the sample was blotted, washed in three drops of water and then stained with 2% (weight/vol) uranyl acetate. Images were recorded on a Gatan UltraScan4000 CCD camera (Gatan) on a Tecnai T12 BioTWINLaB6 electron microscope operating at a voltage of 120 kV.

For cryo-EM, the ternary complex was prepared by incubating 100 μM SUVs, 10 μM SepF and 10 μM FtsZ in polymerization buffer (100 mM KCl, 10 mM MgCl_2_, 25 mM Pipes pH 6.9; 3 mM GTP) for 10 minutes at room temperature. 5 µl of sample were deposited onto glow discharged lacey carbon copper grids (Ted Pella) and plunge-frozen in liquid ethane using a Leica EM-CPC. Cryo-EM data acquisition was performed on a JEOL 2200FS (Jeol, Japan) 200 kV cryo-electron microscope equipped with an Omega in-column energy filter. High magnification (30.000 x, corresponding pixel size 0.32 nm) zero-loss (slit: 20 eV) images were collected at nominal defocus between 1 and 4 µm depending on the experiment on a Gatan USC1000 slow scan CCD camera.

### Mass spectrometry

Strains expressing Scarlet, SepF-Scarlet and SepF_K125E/F131A_-Scarlet were grown in CGXII minimal media supplemented with 4% sucrose and 1% gluconate for 6 hrs at 30 °C. Cells were harvested, washed and normalized by resuspending cell pellets in PBS-T (1X PBS, 0.1% v/v Tween-80) to give a final OD_600_ of 3. The cell suspensions were cross-linked with 0.25% v/v of formaldehyde for 20 min at 30 °C with gentle agitation. The crosslinking reaction was stopped by adding 1.25 M glycine and incubated for 5 min at room temperature. Cells were resuspended in 50 mM Bis-Tris pH 7.4, 75 mM aminocaproic acid, 1 mM MgSO4, 1x Benzonase, 1x Complete Protease inhibitors cocktail (ROCHE) and disrupted at 4 °C with 0.1 mm glass beads and using a PRECELLYS 24 homogenizer. After lysis, 0.1 % DDM was added and the total extract was incubated for 1h at 4 °C. Cell lysates were centrifuged at 4 °C for 10 min at 14 000 rpm and the soluble fraction was collected and protein levels quantified.

Detergent-solubilized protein extracts were incubated with 50 μl of magnetic beads crosslinked with 10 μg of purified anti-Scarlet antibody (produced by Covalab). Beads were collected with a magnetic stand and washed 3 times with lysis buffer and bound material was eluted in 0.1 M glycine pH 2.5 for 10 min at 4 °C, and immediately neutralized by adding 1M Tris pH 9. The protein samples were reduced, denatured (2M Urea) and alkylated with iodoacetamide prior to treatment with trypsin. Tryptic digests were cleaned on a POROS^TM^ R2 resin (Thermo Fisher), vacuum dried and resuspended in phase A buffer (0.1% formic acid). Tryptic peptides from three replicates of each condition were analysed using a nano-HPLC (Ultimate 3000, Thermo) coupled to a hybrid quadrupole-orbitrap mass spectrometer (Q-Exactive Plus, Thermo). Peptide mixtures were separated on a C18 column (PepMap® RSLC, 0.075×500 mm, 2 μm, 100 Ǻ) using a 65 min gradient of mobile phase B from 0% to 55% (A: 0.1% formic acid; B: 0.1% formic acid in acetonitrile). Online MS analysis was carried out in a data dependent mode (MS followed by MS/MS of the top 12 ions) using a dynamic exclusion list. PatternLab for Proteomics software^67^ was used for protein identification against a target-reverse *C. glutamicum* database (Uniprot November 2018) to which the sequences of Scarlet, SepF-Scarlet and -SepF_K125E/F131A_-Scarlet were added.

Patternlab for proteomics was used for label-free quantitation analyses using extracted ion chromatogram (XIC). To calculate fold enrichment of FtsZ in pull down analyses, FtsZ signals (ΣXIC signal of detected peptides in each replicate) were normalized by comparing the ratio FtsZ/SepF in each case (for that we considered the intensity of SepF as (SepF-Scarlet minus Scarlet) or (SepF_K125E/F131A_-Scarlet minus Scarlet). The results are shown in Supplementary Table 2.

### Phylogenetic analysis

We performed sequence homology searches with HMMER ^68^ using SEPF_BACSU/ Q8NNN6_CORGL as queries against the UniprotKB database. More than 4000 sequences were then clustered at 76% with CD-HIT, and after removal of short (<120 residues) and long (>240) candidates, 1800 sequences were kept. Redundancy was further reduced at 50% sequence identity with clustering programs CD-HIT ^69^ and MMseqs2 ^70^ separately, merging both outputs and removing duplicate sequences. We used this non-redundant dataset of 839 sequences with wide taxonomic distribution to compute a multiple alignment with MAFFT ^71^ (l-insi option). Before phylogenetic reconstruction, we used trimAL ^72^ to remove columns with >80% gaps. Finally, we built the SepF tree with PhyML 3.3 ^73^ using the LG+CAT substitution model, chosen with SMS^74^. Trees were visualised using FigTree v1.4.3.

## Acknowledgements

We thank A. Ducret for help with MicrobeJ, F. Gubellini for help with electron microscopy, M. Bott and M. Baumgart for the pk19-P3323-*lcpA* plasmid and help with corynebacterial genetics, and H. Gramajo for the pTGR5 plasmid. We gratefully acknowledge the core facilities at the Institut Pasteur C2RT, in particular G. Pehau-Arnaudet (UBI), B. Raynal, S. Brule (PFBMI), P. Weber, C. Pissis (PFC), and J. Fernandes (UtechS PBI / Imagopole, supported by France BioImaging; ANR-10–INSB–04; Investments for the Future). We thank the staff of ESRF and of EMBL-Grenoble for assistance and support in using beamlines ID30B and ID23-1, and the staff of SOLEIL Synchrotron for assistance in using the beamline Disco. We acknowledge the PICT-IBISA for providing access to the cryo-EM facility at Orsay.

This work was partially supported by grants from the Institut Pasteur (Paris), the CNRS (France) and the Agence Nationale de la Recherche (PhoCellDiv, ANR-18-CE11-0017-01). A.S. is part of the Pasteur - Paris University (PPU) International PhD Program, funded by the European Union’s Horizon 2020 research and innovation programme under the Marie Sklodowska-Curie grant agreement No 665807. Q.G. was funded by MTCI PhD school (ED 563); A.V. was supported by a DIM MalInf (infectious diseases) grant. M.G. acknowledges support from Programa de Desarrollo de las Ciencias Básicas and Sistema Nacional de Investigación e Innovación, Uruguay.

## Author contributions

A.S., M.M., Q.G., M.B.A. and A.M.W. conducted the protein biochemistry, cell biology and genetic experiments, and purified proteins for structural and biophysical studies. P.E. and A.S. carried out the biochemical and biophysical studies of protein-protein interactions. A.V, A.C. and A.S. carried out binding studies of lipid membrane-protein interactions. A.S., A.H., A.M.W. and P.M.A. carried out the crystallogenesis and crystallographic studies. M.G. and A.M.W. performed the phylogeny analyses. M.V.N. contributed essential reagents (fluorescent labelled amino acids). A.S., S.T. and A.M.W. performed the cryo-EM and negative stain EM studies. R.D. carried out the mass spectrometry work. A.M.W. and P.M.A. wrote the paper. All authors edited the paper.

## Competing interests

The authors declare no competing financial interests.

## References

1. Bi, E. F. & Lutkenhaus, J. FtsZ ring structure associated with division in *Escherichia coli*. Nature 354, 161–164 (1991).

2. Du, S. & Lutkenhaus, J. At the Heart of Bacterial Cytokinesis: The Z Ring. Trends Microbiol. 1–11 (2019).

3. Levin, P. A., Kurtser, I. G. & Grossman, A. D. Identification and characterization of a negative regulator of FtsZ ring formation in *Bacillus subtilis*. Proc. Natl. Acad. Sci. USA 96, 9642–9647 (1999).

4. Gueiros-Filho, F. J. & Losick, R. A widely conserved bacterial cell division protein that promotes assembly of the tubulin-like protein FtsZ. Genes Dev. 16, 2544–2556 (2002).

5. Ebersbach, G., Galli, E., Møller-Jensen, J., Löwe, J. & Gerdes, K. Novel coiled-coil cell division factor ZapB stimulates Z ring assembly and cell division. Mol. Microbiol. 68, 720–735 (2008).

6. Durand-Heredia, J. M., Yu, H. H., De Carlo, S., Lesser, C. F. & Janakiraman, A. Identification and characterization of ZapC, a stabilizer of the FtsZ ring in *Escherichia coli*. J. Bacteriol. 193, 1405–1413 (2011).

7. Durand-Heredia, J., Rivkin, E., Fan, G., Morales, J. & Janakiraman, A. Identification of ZapD as a cell division factor that promotes the assembly of FtsZ in *Escherichia coli*. J. Bacteriol. 194, 3189–3198 (2012).

8. Pichoff, S. & Lutkenhaus, J. Tethering the Z ring to the membrane through a conserved membrane targeting sequence in FtsA. Mol. Microbiol. 55, 1722–1734 (2005).

9. Hale, C. A. & de Boer, P. A. Direct binding of FtsZ to ZipA, an essential component of the septal ring structure that mediates cell division in *E. coli*. Cell 88, 175–185 (1997).

10. Loose, M. & Mitchison, T. J. The bacterial cell division proteins FtsA and FtsZ self-organize into dynamic cytoskeletal patterns. Nat. Cell Biol. 16, 38–46 (2014).

11. Duman, R. et al. Structural and genetic analyses reveal the protein SepF as a new membrane anchor for the Z ring. Proc. Natl. Acad. Sci. USA 110, E4601–10 (2013).

12. Bernhardt, T. G. & de Boer, P. A. J. SlmA, a Nucleoid-Associated, FtsZ Binding Protein Required for Blocking Septal Ring Assembly over Chromosomes in *E. coli*. Mol. Cell 18, 555–564 (2005).

13. Wu, L. J. & Errington, J. Coordination of cell division and chromosome segregation by a nucleoid occlusion protein in *Bacillus subtilis*. Cell 117, 915–925 (2004).

14. Shen, B. & Lutkenhaus, J. The conserved C-terminal tail of FtsZ is required for the septal localization and division inhibitory activity of MinC(C)/MinD. Mol. Microbiol. 72, 410–424 (2009).

15. Ma, X. & Margolin, W. Genetic and functional analyses of the conserved C-terminal core domain of *Escherichia coli* FtsZ. J. Bacteriol. 181, 7531–7544 (1999).

16. Donovan, C. & Bramkamp, M. Cell division in *Corynebacterineae*. Front. Microbiol 5, 132 (2014).

17. Letek, M. et al. Cell growth and cell division in the rod-shaped actinomycete *Corynebacterium glutamicum*. Antonie Van Leeuwenhoek 94, 99–109 (2008).

18. Ishikawa, S., Kawai, Y., Hiramatsu, K., Kuwano, M. & Ogasawara, N. A new FtsZ-interacting protein, YlmF, complements the activity of FtsA during progression of cell division in *Bacillus subtilis*. Mol. Microbiol. 60, 1364–1380 (2006).

19. Hamoen, L. W., Meile, J.-C., de Jong, W., Noirot, P. & Errington, J. SepF, a novel FtsZ-interacting protein required for a late step in cell division. Mol. Microbiol. 59, 989–999 (2006).

20. Gola, S., Munder, T., Casonato, S., Manganelli, R. & Vicente, M. The essential role of SepF in mycobacterial division. Mol. Microbiol. 97, 560–576 (2015).

21. Marbouty, M., Saguez, C., Cassier-Chauvat, C. & Chauvat, F. Characterization of the FtsZ-interacting septal proteins SepF and Ftn6 in the spherical-celled cyanobacterium *Synechocystis* strain PCC 6803. J. Bacteriol. 191, 6178–6185 (2009).

22. Gündoğdu, M. E. et al. Large ring polymers align FtsZ polymers for normal septum formation. EMBO J. 30, 617–626 (2011).

23. Krupka, M. et al. Role of the FtsA C terminus as a switch for polymerization and membrane association. mBio 5, e02221 (2014).

24. Krupka, M. et al. *Escherichia coli* FtsA forms lipid-bound minirings that antagonize lateral interactions between FtsZ protofilaments. Nat. Commun. 8, 1–12 (2017).

25. Conti, J., Viola, M. G. & Camberg, J. L. FtsA reshapes membrane architecture and remodels the Z-ring in *Escherichia coli*. Mol. Microbiol. 107, 558–576 (2018).

26. Honrubia, M.P., Ramos, A. & Gil, J.A. The cell division genes ftsQ and ftsZ, but not the three downstream open reading frames YFIH, ORF5 and ORF6, are essential for growth and viability in *Brevibacterium lactofermentum* ATCC 13869. Mol. Genet. Genomics 265, 1022–1030 (2001).

27. Baumgart, M. et al. IpsA, a novel LacI-type regulator, is required for inositol-derived lipid formation in Corynebacteria and Mycobacteria. BMC Biol. 11, 122 (2013).

28. Pfeifer-Sancar, K., Mentz, A., Rückert, C. & Kalinowski, J. Comprehensive analysis of the *Corynebacterium glutamicum* transcriptome using an improved RNAseq technique. BMC Genomics 14, 888 (2013).

29. Kuru, E., Tekkam, S., Hall, E., Brun, Y. V. & Van Nieuwenhze, M. S. Synthesis of fluorescent D-amino acids and their use for probing peptidoglycan synthesis and bacterial growth in situ. Nat. Protoc. 10, 33–52 (2014).

30. Frunzke, J., Engels, V., Hasenbein, S., Gätgens, C. & Bott, M. Coordinated regulation of gluconate catabolism and glucose uptake in *Corynebacterium glutamicum* by two functionally equivalent transcriptional regulators, GntR1 and GntR2. Mol. Microbiol. 67, 305–322 (2007).

31. Singh, J. K., Makde, R. D., Kumar, V. & Panda, D. SepF increases the assembly and bundling of FtsZ polymers and stabilizes FtsZ protofilaments by binding along its length. J. Biol. Chem. 283, 31116–31124 (2008).

32. Mosyak, L. et al. The bacterial cell-division protein ZipA and its interaction with an FtsZ fragment revealed by X-ray crystallography. EMBO J. 19, 3179–3191 (2000).

33. Szwedziak, P., Wang, Q., Freund, S. M. & Löwe, J. FtsA forms actin-like protofilaments. EMBO J. 31, 2249–2260 (2012).

34. Schumacher, M. A., Huang, K.-H., Zeng, W. & Janakiraman, A. Structure of the Z ring-associated protein, ZapD, bound to the C-terminal domain of the tubulin-like protein, FtsZ, suggests mechanism of Z ring stabilization through FtsZ cross-linking. J. Biol. Chem. 292, 3740–3750 (2017).

35. Huang, K.-H., Durand-Heredia, J. & Janakiraman, A. FtsZ ring stability: of bundles, tubules, crosslinks, and curves. J. Bacteriol. 195, 1859–1868 (2013).

36. Schumacher, M. A. & Zeng, W. Structures of the nucleoid occlusion protein SlmA bound to DNA and the C-terminal domain of the cytoskeletal protein FtsZ. Proc. Natl. Acad. Sci. USA 113, 4988–4993 (2016).

37. Addinall, S. G. & Lutkenhaus, J. FtsA is localized to the septum in an FtsZ-dependent manner. J. Bacteriol. 178, 7167–7172 (1996).

38. Bisson-Filho, A. W. et al. Treadmilling by FtsZ filaments drives peptidoglycan synthesis and bacterial cell division. Science 355, 739–743 (2017).

39. Yang, X. et al. GTPase activity-coupled treadmilling of the bacterial tubulin FtsZ organizes septal cell wall synthesis. Science 355, 744–747 (2017).

40. Osawa, M., Anderson, D. E. & Erickson, H. P. Reconstitution of contractile FtsZ rings in liposomes. Science 320, 792–794 (2008).

41. Ramirez-Diaz, D. A. et al. Treadmilling analysis reveals new insights into dynamic FtsZ ring architecture. PLoS Biol. 16, e2004845–20 (2018).

42. Ramirez-Diaz, D. A., Merino-Salomon, A., Heymann, M. & Schwille, P. Bidirectional FtsZ filament treadmilling promotes membrane constriction via torsional stress. bioRxiv 3, e04601–9 (2019).

43. Krupka, M., Sobrinos-Sanguino, M., Jiménez, M., Rivas, G. & Margolin, W. *Escherichia coli* ZipA organizes FtsZ polymers into dynamic ring-like protofilament structures. mBio 9, e01008–18–15 (2018).

44. Caldas, P. et al. ZapA stabilizes FtsZ filament bundles without slowing down treadmilling dynamics. bioRxiv 1–33 (2019).

45. Rivas, G. & Minton, A. P. Macromolecular crowding in vitro, in vivo, and in between. Trends Biochem. Sci. 41, 970–981 (2016).

46. van den Berg, J., Boersma, A. J. & Poolman, B. Microorganisms maintain crowding homeostasis. Nat. Rev. Microbiol. 15, 309–318 (2017).

47. Xiao, J. & Goley, E. D. Redefining the roles of the FtsZ-ring in bacterial cytokinesis. Curr. Opin. Microbiol. 34, 90–96 (2016).

48. Keilhauer, C., Eggeling, L. & Sahm, H. Isoleucine synthesis in *Corynebacterium glutamicum*: molecular analysis of the ilvB-ilvN-ilvC operon. J. Bacteriol. 175, 5595–5603 (1993).

49. Ravasi, P., Peiru, S., Gramajo, H. & Menzella, H. G. Design and testing of a synthetic biology framework for genetic engineering of *Corynebacterium glutamicum*. Microb. Cell Fact. 11, 1–1 (2012).

50. Lausberg, F., Chattopadhyay, A. R., Heyer, A., Eggeling, L. & Freudl, R. A tetracycline inducible expression vector for *Corynebacterium glutamicum* allowing tightly regulable gene expression. Plasmid 68, 142–147 (2012).

51. Bindels, D. S. et al. mScarlet: a bright monomeric red fluorescent protein for cellular imaging. Nat. Methods 14, 53–56 (2016).

52. Shaner, N. C. et al. A bright monomeric green fluorescent protein derived from *Branchiostoma lanceolatum*. Nat. Methods 10, 407–409 (2013).

53. Baumgart, M., Schubert, K., Bramkamp, M. & Frunzke, J. Impact of LytR-CpsA-Psr proteins on cell wall biosynthesis in *Corynebacterium glutamicum*. J. Bacteriol. 198, 3045–3059 (2016).

54. Studier, F. W. Protein production by auto-induction in high density shaking cultures. Protein Expr. Purif. 41, 207–234 (2005).

55. Kabsch, W. XDS. Acta Crystallogr. D66, 125–132 (2010).

56. Winn, M. D. et al. Overview of the CCP4 suite and current developments. Acta Cryst. D67, 235–242 1–8 (2011).

57. McCoy, A. J. et al. Phaser crystallographic software. J Appl. Crystallogr. 40, 658–674 (2007).

58. Schneider, T. R. & Sheldrick, G. M. Substructure solution with SHELXD. Acta Cryst. D58, 1772–1779 (2002).

59. Cowtan, K. The Buccaneer software for automated model building. 1. Tracing protein chains. Acta Cryst. D62, 1002–1011 (2006).

60. Emsley, P. & Cowtan, K. Coot: model-building tools for molecular graphics. Acta Cryst. D60, 2126–2132 (2004).

61. Bricogne, G. et al. Buster version 2.10.3. Cambridge, United Kingdom: Global Phasing Ltd. (2017).

62. Pettersen, E. F. et al. UCSF Chimera: a visualization system for exploratory research and analysis. J. Comput. Chem. 25, 1605–1612 (2004).

63. Micsonai, A. et al. Accurate secondary structure prediction and fold recognition for circular dichroism spectroscopy. Proc. Natl. Acad. Sci. USA 112, E3095–E3103 (2015).

64. Jomain, J.-B. et al. Structural and thermodynamic bases for the design of pure prolactin receptor antagonists: X-ray structure of Del1-9-G129R-hPRL. J. Biol. Chem. 282, 33118–33131 (2007).

65. Schindelin, J. et al. Fiji: an open-source platform for biological-image analysis. Nat Methods 9, 676–682 (2012).

66. Ducret, A., Quardokus, E. M. & Brun, Y. V. MicrobeJ, a tool for high throughput bacterial cell detection and quantitative analysis. Nat. Microbiol. 1, 1–7 (2016).

67. Carvalho, P. C. et al. Integrated analysis of shotgun proteomic data with PatternLab for proteomics 4.0. Nat Protoc 11, 102–117 (2015).

68. Eddy, S. R. Profile hidden Markov models. Bioinformatics 14, 755–763 (1998).

69. Li, W. & Godzik, A. Cd-hit: a fast program for clustering and comparing large sets of protein or nucleotide sequences. Bioinformatics 22, 1658–1659 (2006).

70. Steinegger, M. & Söding, J. MMseqs2 enables sensitive protein sequence searching for the analysis of massive data sets. Nat. Biotechnol. 35, 1026–1028 (2017).

71. Katoh, K. & Standley, D. M. MAFFT: iterative refinement and additional methods. Methods Mol. Biol. 1079, 131–146 (2013).

72. Capella-Gutierrez, S., Silla-Martinez, J. M. & Gabaldon, T. TrimAl: a tool for automated alignment trimming in large-scale phylogenetic analyses. Bioinformatics 25, 1972–1973 (2009).

73. Guindon, S. et al. New algorithms and methods to estimate maximum-likelihood phylogenies: assessing the performance of PhyML 3.0. Systematic Biol. 59, 307–321 (2010).

74. Lefort, V., Longueville, J.-E. & Gascuel, O. SMS: Smart model selection in PhyML. Mol. Biol. Evol. 34, 2422–2424 (2017).

