## Supplemental Material for "Essential dynamic interdependence of FtsZ and SepF for Z-ring and septum formation in *Corynebacterium glutamicum*"

**Supplementary Methods, Supplementary Figures 1-22 and Supplementary Tables 1-5.**

### Supplementary Methods

#### Dynamic light Scattering:

Before DLS experiments, protein samples were centrifuged at 25.000 x g for 15min at 4°C. SUVs with or without SepF (or SepF mutants) were used at a final concentration of 50 µM each in 100 mM KCl, 10 mM $MgCl_2$, and 25 mM Pipes pH 6.9 Buffer and the reaction was carried out for 10 minutes at room-temperature. Polydispersity of the samples was measured using Dyna-pro Plate Reader (Wyatt technology) equipped with an 830 nm laser and a temperature control module. The *Dynamics* software (version 7.9) was used to schedule data acquisition and data analysis. For each well, 20 measurements of 10 seconds were averaged and this operation was repeated 20 times for each condition at 25°C. For samples containing $FtsZ_{CTD}$, the peptide was used at a final concentration of 100 µM. All the experiments were carried out as minimum 3 biological and technical replicates.

#### Analytical ultracentrifugation (AUC):

Protein samples were desalted using a PD Minitrap G-25 column (Sigma-Aldrich) into AUC buffer (50 mM Hepes pH 7.4, 150 mM KCl, 10 mM MgCl2) and analyzed at 1-2 mg/ml concentration. Samples were centrifuged at 42000 rpm in a Beckman Coulter XL-1 analytical ultracentrifuge at 20 °C in a four-hole AN 60–Ti rotor equipped with 12-mm double-sector epoxy centerpieces. Detection of SepF concentration as a function of radial position and time was performed by optical density measurements at 280 nm and interferometry. Data analysis for sedimentation velocity was performed by continuous size distribution c(s) using Sedfit software version 15.01.

**References**

1. Hanahan, D. Studies on transformation of *Escherichia coli* with plasmids. *J. Mol. Biol.* **166,** 557–580 (1983).
2. Studier, F. W. & Moffatt, B. A. Use of bacteriophage T7 RNA polymerase to direct selective high-level expression of cloned genes. *J. Mol. Biol.* **189,** 113–130 (1986).
3. Haskins, D. *Epicentre Forum* **11**, 6 (2004).
4. Kinoshita, S. & Udaka, S. Studies on the amino acid fermentation. *J. Gen. Appl. Microbiol.* **3,** 193–205 (1957).
5. Schäfer, A. *et al.* Small mobilizable multi-purpose cloning vectors derived from the *Escherichia coli* plasmids pK18 and pK19: selection of defined deletions in the chromosome of *Corynebacterium glutamicum*. *Gene* **145,** 69–73 (1994).
6. Baumgart, M., Schubert, K., Bramkamp, M. & Frunzke, J. Impact of LytR-CpsA-Psr Proteins on Cell Wall Biosynthesis in *Corynebacterium glutamicum*. *J. Bacteriol.* **198,** 3045–3059 (2016).
7. Ravasi, P., Peiru, S., Gramajo, H. & Menzella, H. G. Design and testing of a synthetic biology framework for genetic engineering of *Corynebacterium glutamicum*. *Microb. Cell Fact.* **11,** 147 (2012).
8. Lausberg, F., Chattopadhyay, A. R., Heyer, A., Eggeling, L. & Freudl, R. A tetracycline inducible expression vector for *Corynebacterium glutamicum* allowing tightly regulable gene expression. *Plasmid* **68,** 142–147 (2012).
9. Laskowski, R. A. & Swindells, M. B. LigPlot+: Multiple ligand-protein interaction diagrams for drug discovery. *J. Chem. Inf. Model* **51**, 2778-2786 (2011).

### Supplementary Figures

**Supplementary Figure 1: Complementation of the *$P_{ino}$-sepF* strain during SepF depletion by the expression of *$P_{tet}$-sepF* upon tetracycline addition. a.** Growth curves comparing WT-$P_{tet}$ (blue) with $P_{ino}$-*sepF*-$P_{tet}$-*sepF* (red) in 1% *myo*-inositol + 50 ng/ml tetracycline. **b.** Western blot of whole cell extracts corresponding to the above growth curve at different time points (0h, 3h, 6h, 9h, and over-night (ON)) probed with anti-SepF antibodies to confirm SepF expression during complementation of the SepF depleted strain. **c.** Violin plot of triplicate analysis showing the distribution of cell length at time point 4.5 hours after *myo*-inositol and tetracycline addition for WT-$P_{tet}$ (blue) and $P_{ino}$-*sepF*-$P_{tet}$-*sepF* (orange). Medians and standard deviations of cell lengths are shown in Table T5.

a

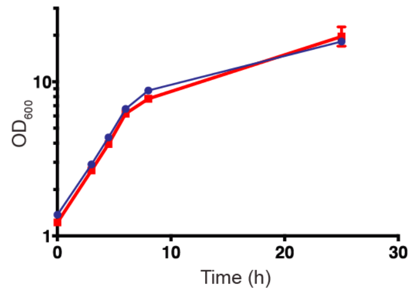

b

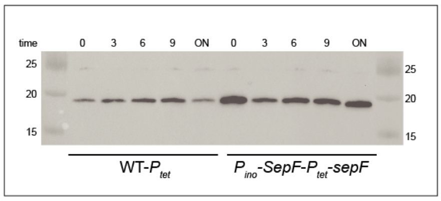

c

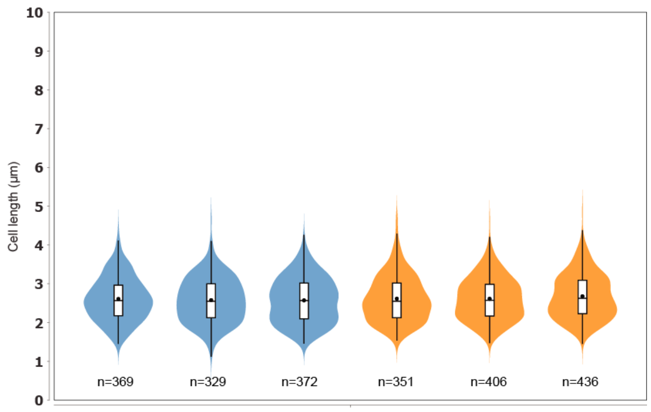

**Supplementary Figure 2: Triplicate analysis of fluorescent HADA localization and cell length during SepF depletion. a.** Heat maps representing the localization pattern of HADA at 0, 3 and 6 hours after *myo*-inositol addition for $P_{ino}$-*sepF* (left group of triplicates) and WT (right group of triplicates) strains. **b.** Violin plot of triplicate analysis showing the distribution of cell length at time points 0, 3, 6 hours after *myo*-inositol addition for $P_{ino}$-*sepF* and WT. Medians and standard deviations of cell lengths are shown in Table T5. Medians between triplicate time points are different (Mann-Whitney) for *Pino-sepF*.

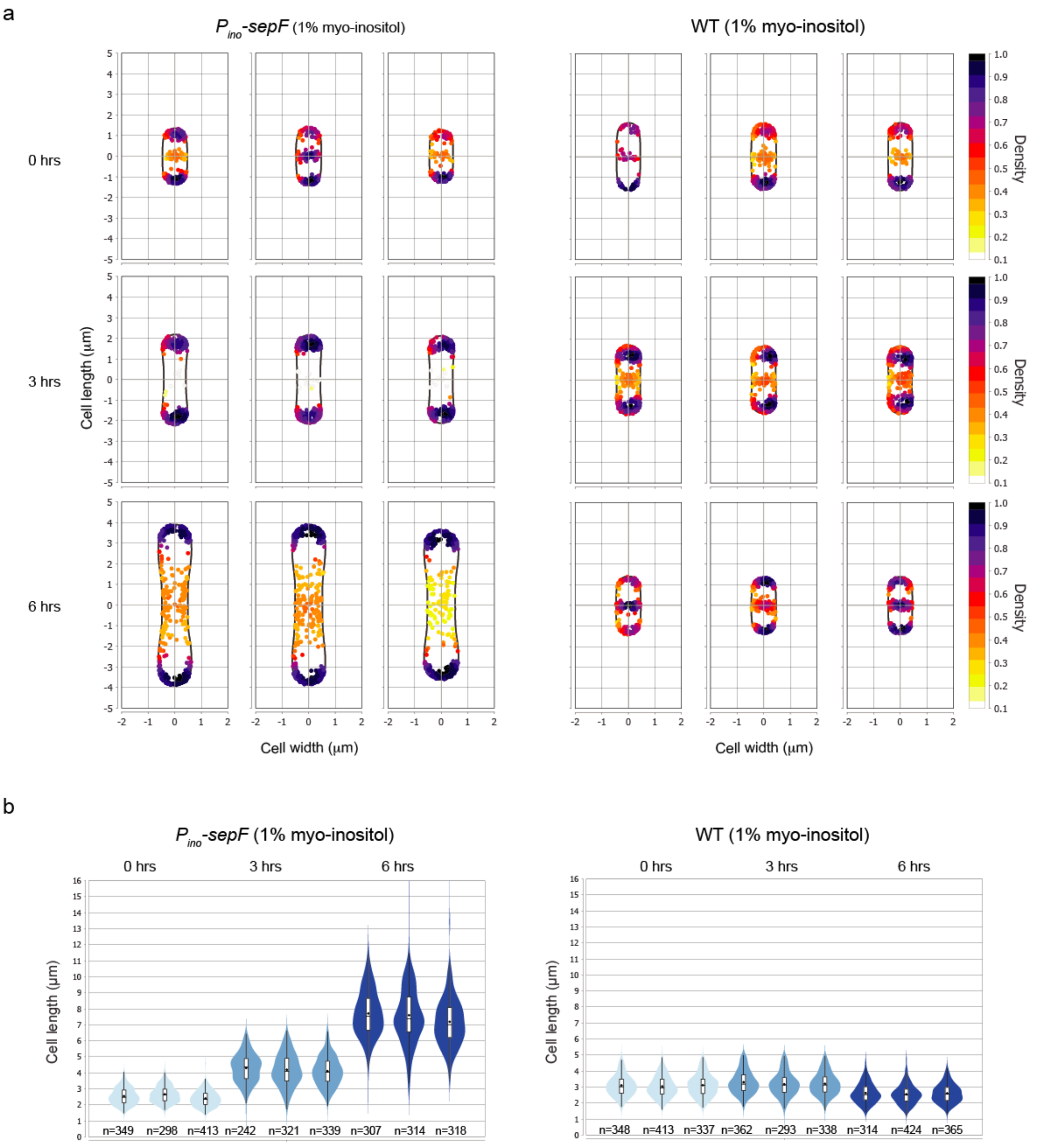

**Supplementary Figure 3: Maximum likelihood phylogenetic tree of bacterial SepF proteins.** Major Phyla are shown in different colors. Grey-shaded areas highlight branches with no detectable *ftsA* homologue, as is the case for *Actinobacteria*, *Cyanobacteria*, *Melainabacteria*, *Tenericutes* and some clades within *Firmicutes*. The scale bar represents average substitutions per site.

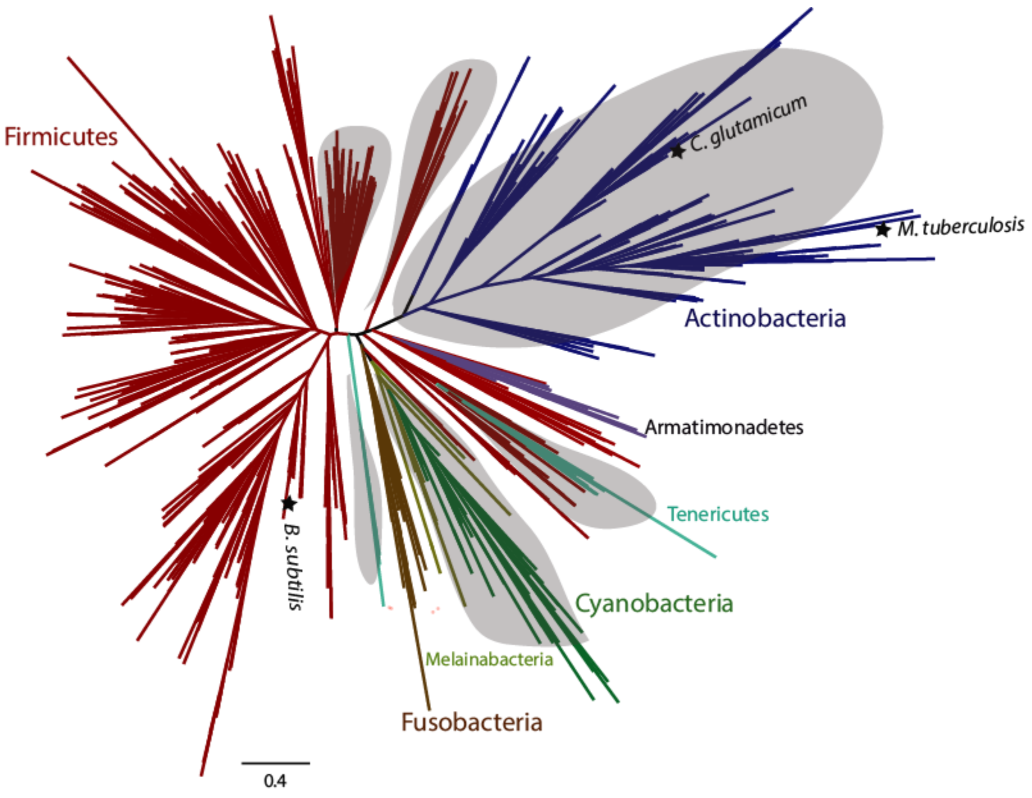

**Supplementary Figure 4: mNeon-FtsZ expression system**: **a.** Schematic representation of the $P_{gntK}$ promotor in the pTGR5 vector backbone. The $P_{gntK}$ promotor is repressed in the presence of 4% sucrose and induced by 1% gluconate. **b.** Western blot of whole cell extracts of WT-$P_{gntK}$-*mneon-ftsZ* cells grown in either 4% sucrose (S) or 4% sucrose + 1% gluconate (G). The anti-FtsZ antibody was used to reveal both endogenous and recombinant FtsZ. **c.** Comparison of the size distribution in exponential phase of WT-$P_{gntK}$ (blue) and WT-$P_{gntK}$-*mneon-ftsZ* (green). Medians and standard deviations of cell lengths are shown in Table T5.

a

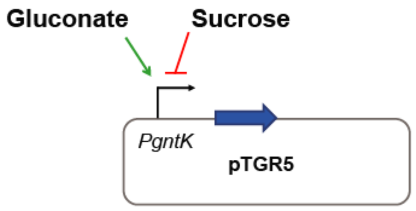

b

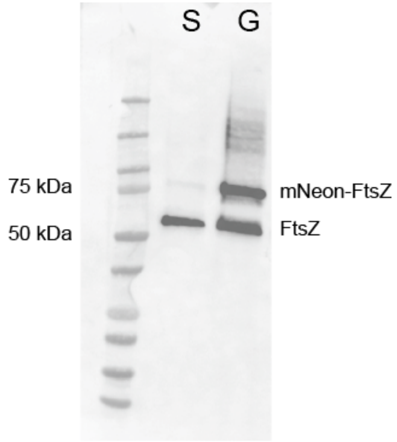

c

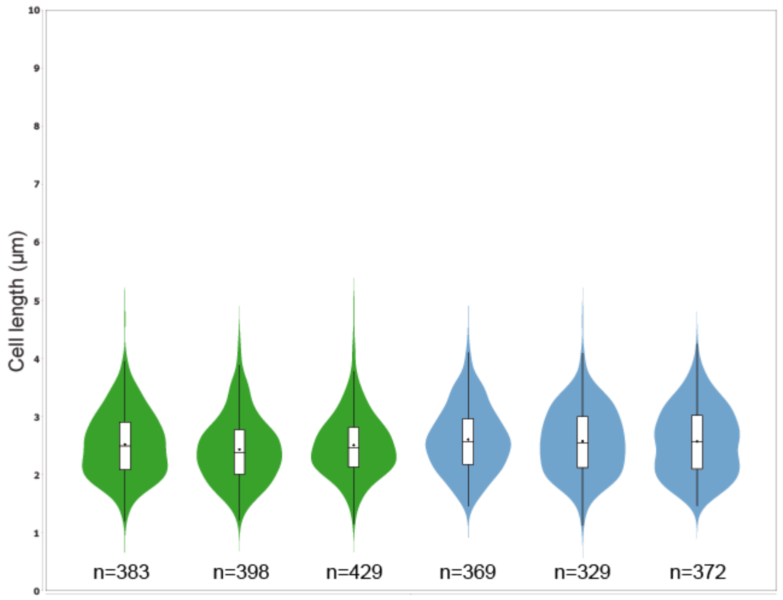

**Supplementary Figure 5: Characterization of the *$P_{ino}$-sepF-$P_{gntK}$-mneon-FtsZ* strain.**

**a.** Violin plot of triplicate analysis showing the cell length distribution at time points 0, 3, 6 hours after *myo*-inositol addition. Medians of triplicates were identical at each time point (Mann-Whitney) and medians of two different time points for one sample were different (Mann-Whitney). **b.** Triplicate heatmaps representing the localization pattern of HADA (left) and mNeon-FtsZ (right) for the above strain at 0, 3 and 6 hours. Medians and standard deviations of cell lengths are shown in Table T5.

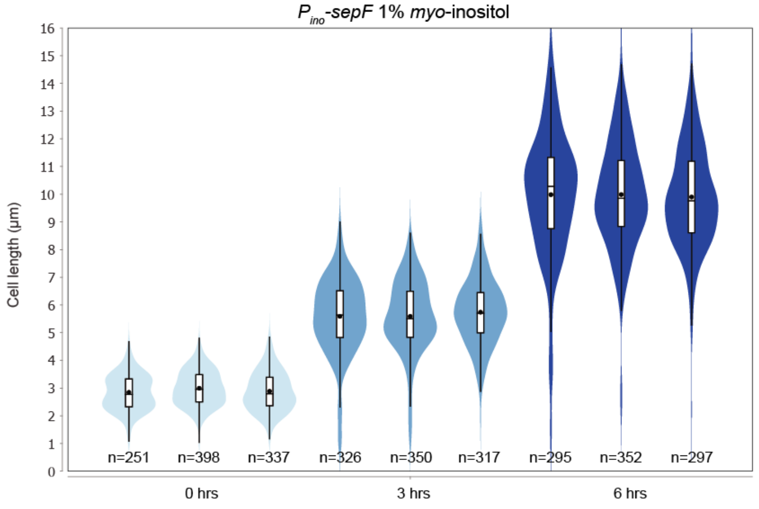

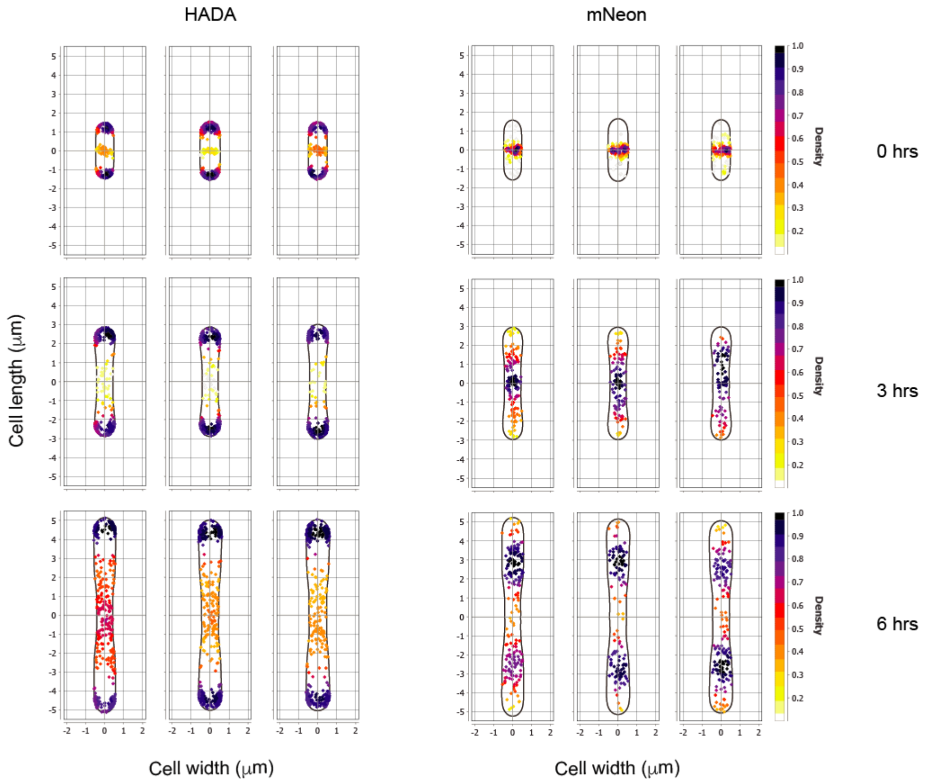

**Supplementary Figure 6. SepF$_M$ peptide-membrane interactions. a.** Amphipathic helix prediction of the membrane-binding region of SepF (SepF$_M$). **b.** Tryptophan fluorescence titration assay using a modified SepF$_M$ peptide (including a C-terminal tryptophan residue) as a function of lipid concentration (see Methods for details). **c.** Circular dichroism spectra of SepF$_M$ in the absence (red) and presence (blue) of SUVs.

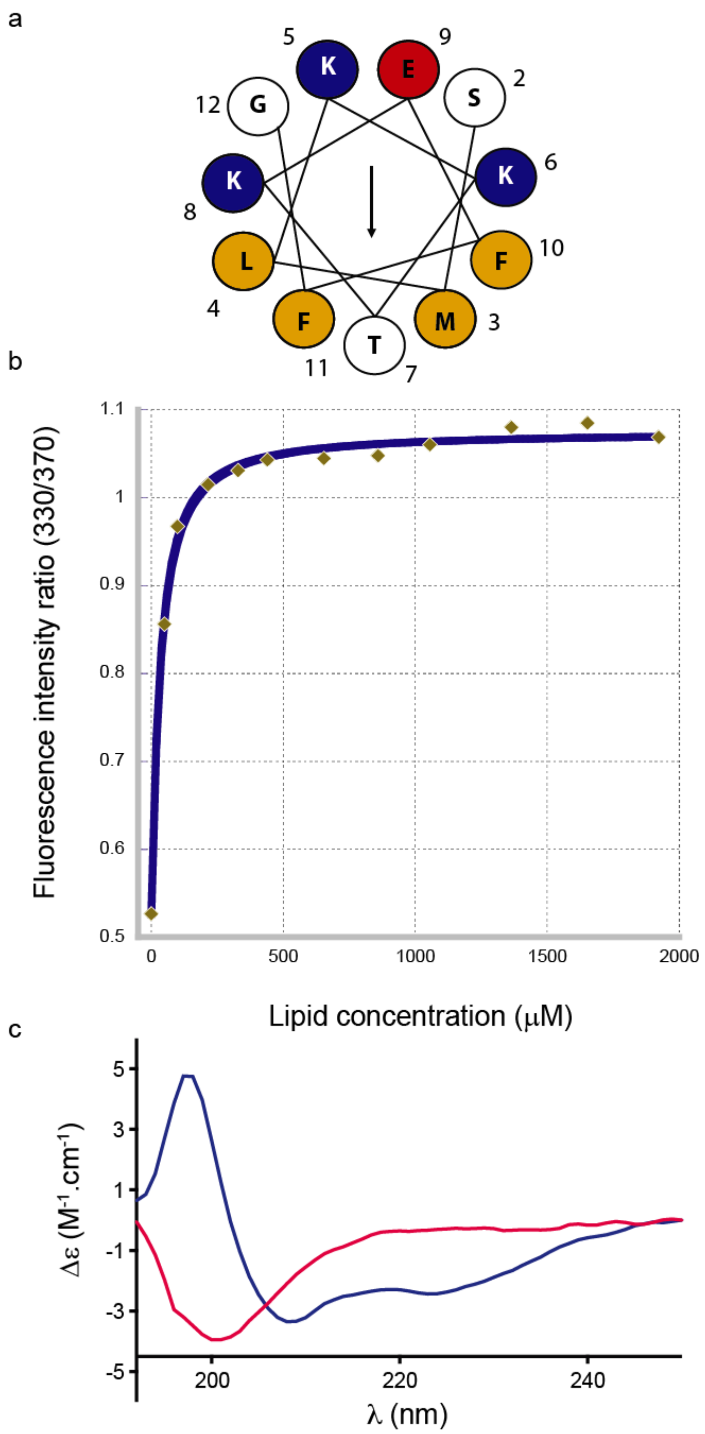

**Supplementary Figure 7: FtsZ polymerization assays in the presence of SepF mutants.** **a.** The absorbance measured at 400 nm for FtsZ alone (15 μM, blue curve) and FtsZ plus SepF$_{\Delta ML}$ (15 μM each, green curve). **b.** The absorbance measured at 400 nm for FtsZ alone (15 μM, blue curve) and FtsZ plus SepF$_{K125E/F131A}$ (15 μM each, red curve).

a b

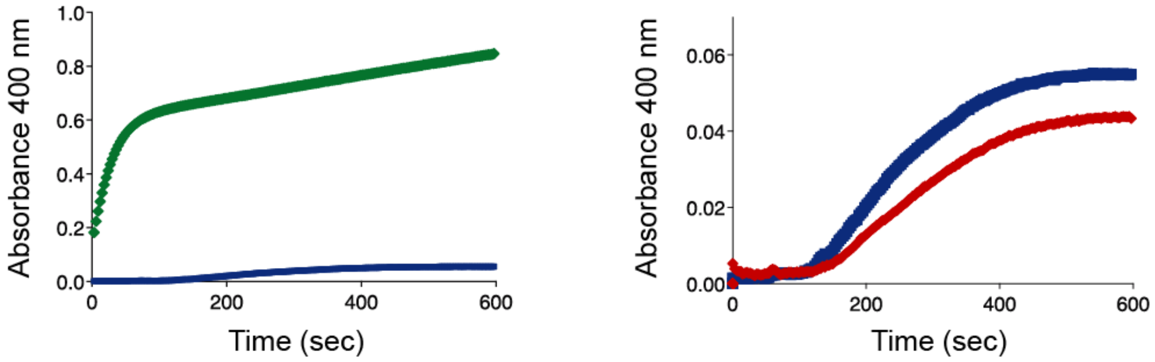

**Supplementary Figure 8: Intermolecular interactions in the SepF-FtsZ$_{CTD}$ complex. a.** Final electron density map of FtsZ$_{CTD}$ contoured at 1.1 σ. **b.** Interactions made by peptide FtsZ$_{CTD}$ (blue) with residues in the SepF binding pocket. The plot was made with LIGPLOT [9].

a b

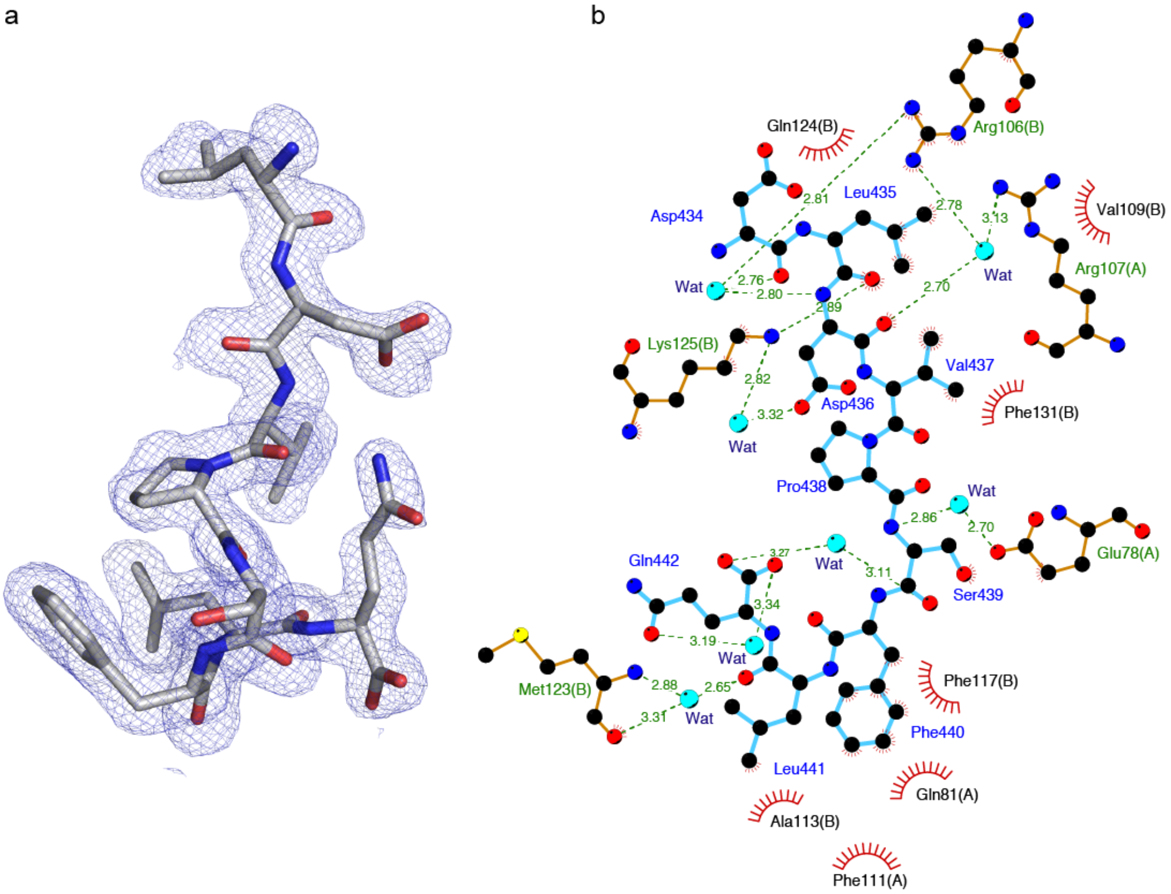

**Supplementary Figure 9: SPR binding profiles and fitted curves for $FtsZ_{CTD}$ interactions with: (a) $SepF_{\Delta ML}$, (b) $SepF_{\Delta ML,F131A}$ and (c) $SepF_{\Delta ML,K125/F131A}$.**

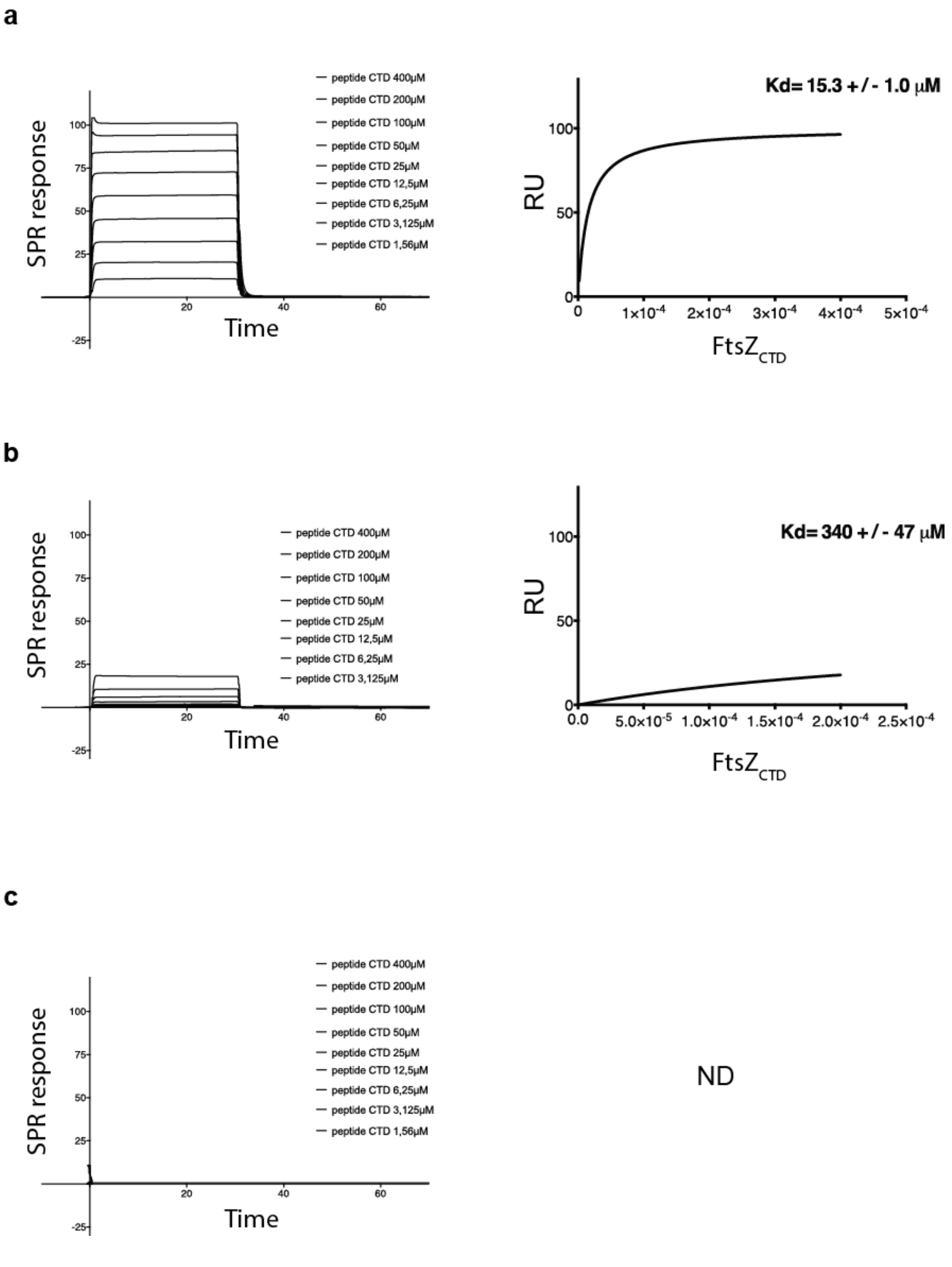

**Supplementary Figure 10: The same conserved hydrophobic residues of $FtsZ_{CTD}$ are involved in protein contacts with SepF and SlmA. a.** Structure of the $FtsZ_{CTD}$ peptide (in stick representation) bound to the *C. glutamicum* SepF binding pocket color-coded according to surface electrostatic potential. **b.** Idem for $FtsZ_{CTD}$ bound to *E. coli* SlmA. **c.** Partial sequence alignment of $FtsZ_{CTD}$ from *C. glutamicum* and *E. coli*. Hydrophobic residues in bold are labelled in **a** and **b**.

a

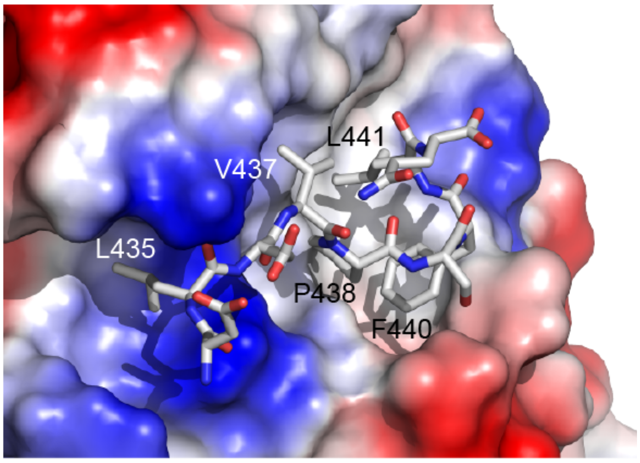

b

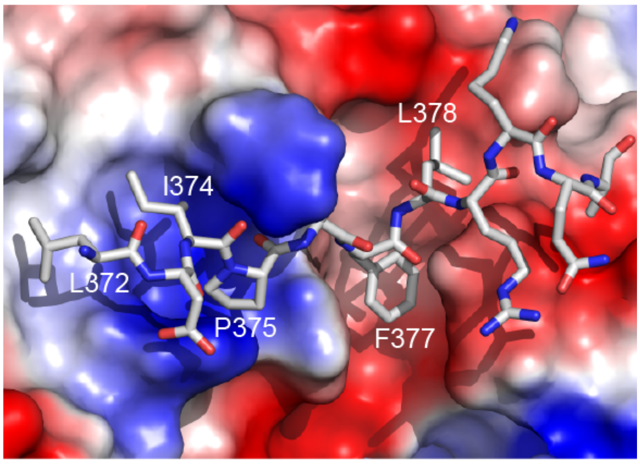

c

| | | | |
|---|---|---|---|
| *E. coli* FtsZ | 371 | YLDIPAFLR | 379 |
| *C. glut* FtsZ | 434 | DLDVPSFLQ | 442 |

**Supplementary Figure 11: Complementation of the $P_{ino}$-*sepF* strain by the expression of $P_{gntK}$-*sepF-scarlet* mutants. a.** Western blot of whole cell extracts corresponding to the growth curve shown in Fig. 3 at different time points (0h, 3h, 6h, ON) probed with anti-SepF antibodies to confirm SepF and mutant expression. **b.** Violin plot of triplicate analysis showing the cell length distribution at 6 hours for the $P_{ino}$-*sepF* strain expressing $P_{gntK}$-*sepF*$_{\Delta ML}$-*scarlet* (blue), $P_{gntK}$-*sepF-scarlet* (green), $P_{gntK}$-*sepF*$_{K125/F131A}$-*scarlet* (red) in 1% *myo*-inositol or $P_{gntK}$ in 0% *myo*-inositol (purple). Medians and standard deviations of cell lengths are shown in Table T5.

a

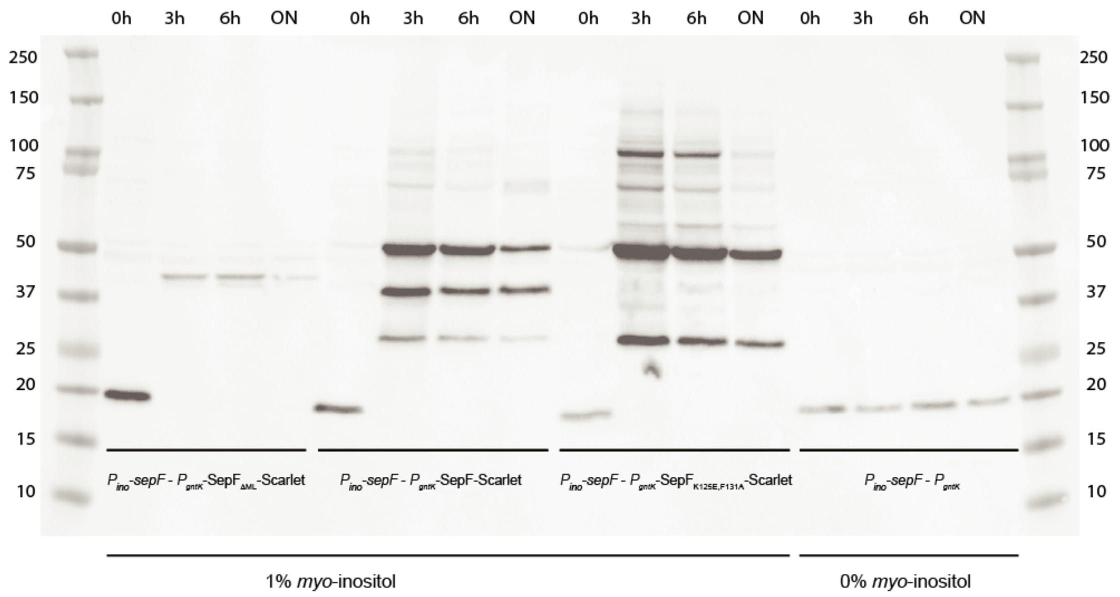

b

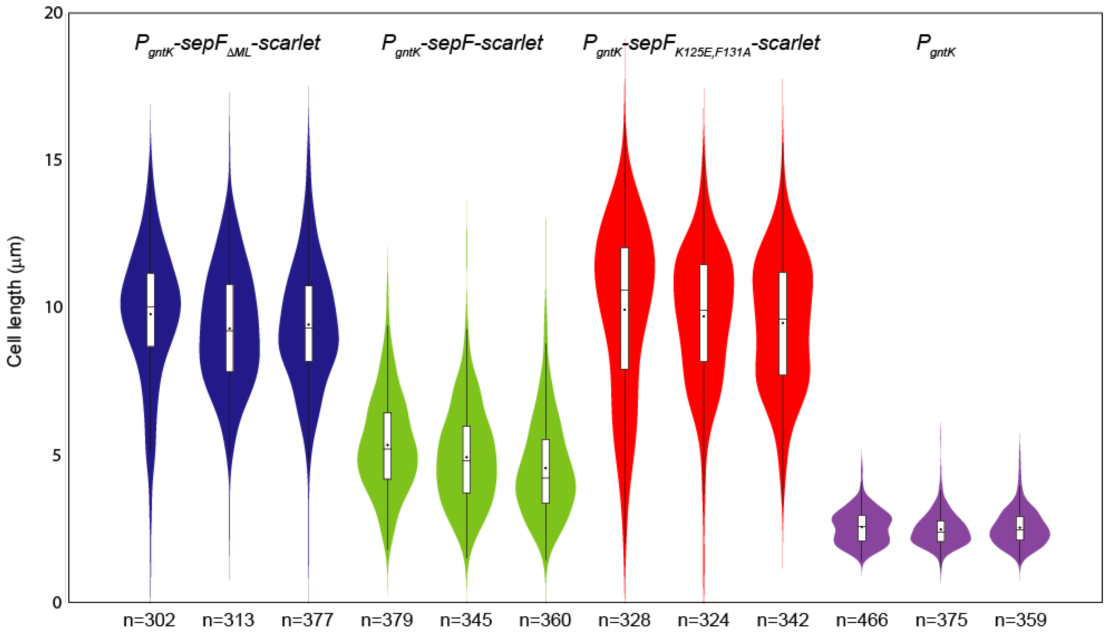

**Supplementary Figure 12: Linear polymers of SepF in the crystal state. a.** Two possible dimerization interfaces identified in the crystal structure of *B. subtilis* SepF (pdb code 3ZIH), mediated respectively by a four-helix bundle (labelled $\alpha$) and by the tight association of opposite sheets (labelled $\beta$), giving rise to linear polymers of the protein. **b-d.** The crystal structures of different constructs of *C. glutamicum* SepF lacking the C-terminal helix $\alpha$3 exhibit a similar molecular arrangement. The panels show the crystal lattices of (**b**) $\text{SepF}_{\Delta ML,\Delta\alpha 3}$-$\text{FtsZ}_{CTD}$, (**c**) $\text{SepF}_{\Delta ML,\Delta\alpha 3}$, and (**d**) PDB code 3P04, projected along two perpendicular directions, showing respectively a lateral (left) and a frontal (right) view of the packing of linear SepF polymers. In each projection, a single polymer is highlighted.

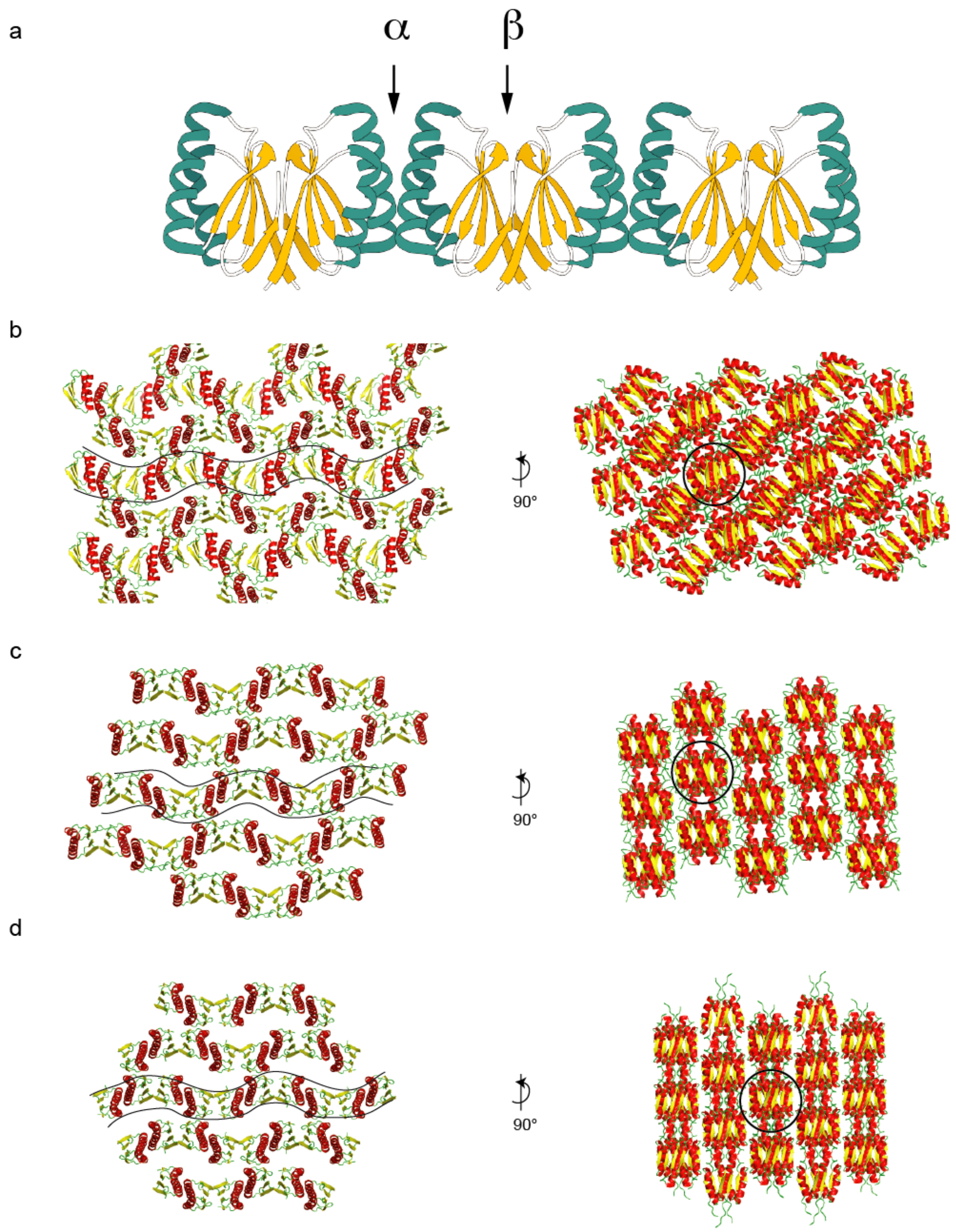

**Supplementary Figure 13: Biophysical characterization of the oligomerization state of SepF and SepF$_{\Delta\alpha3}$. a.** Size exclusion chromatography of SepF (blue) and SepF$_{\Delta\alpha3}$ (red) on a Superdex S200 10/300 column. The molecular weight (MW) markers are shown in gray and the numbers correspond to the MW in kDa. **b.** Analytical ultracentrifugation profile of SepF and SepF$_{\Delta\alpha3}$ detected by interference. For SepF (blue) the main peak showed an estimated molecular weight of 31.3 kDa with a sedimentation coefficient (S) of 2.239 in agreement with the values expected for a SepF dimer. For SepF$_{\Delta\alpha3}$ (green) the main peak showed an estimated molecular weight of 28.8 kDa with a sedimentation coefficient (S) of 2.079 in agreement with the values expected for a SepF$_{\Delta\alpha3}$ dimer.

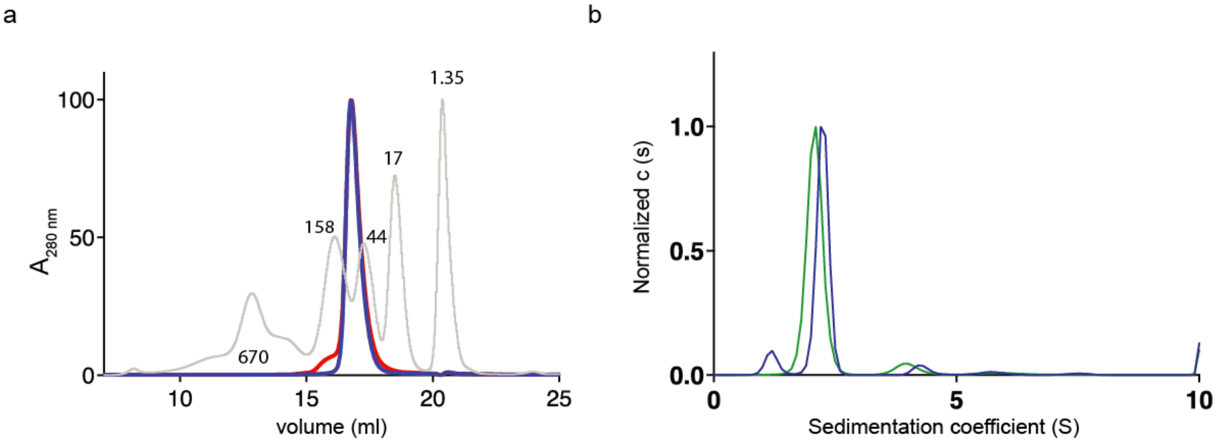

**Supplementary Figure 14: Rings of purified negatively stained *Mtb*SepF$_{\Delta ML}$ (50 µM) observed by electron microscopy.** The scale bar is 50nm.

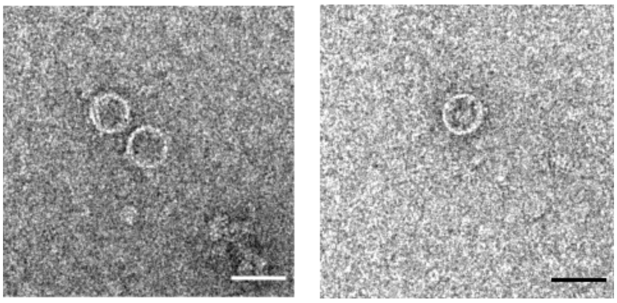

**Supplementary Figure 15: Turbidity assays.** Polymerization of SepF at the indicated protein concentrations was assessed in the presence of lipid membranes (50 μM SUVs).

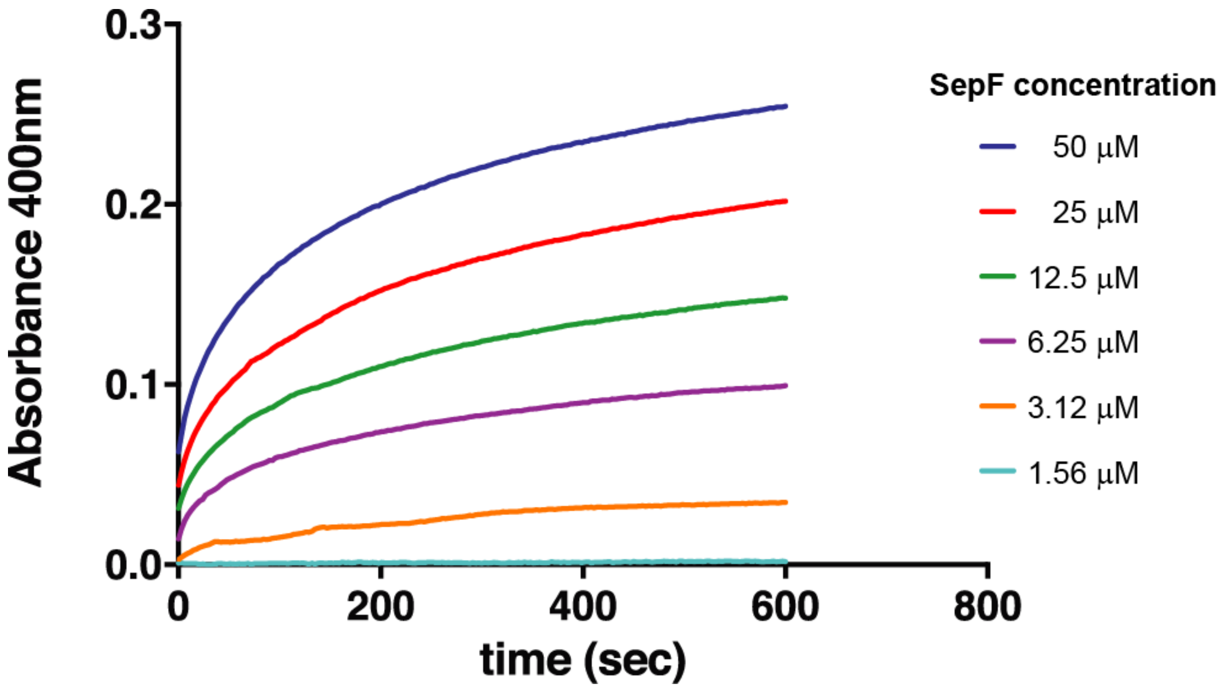

**Supplementary Figure 16: SepF-SUV-FtsZ$_{CTD}$ interaction analyzed by dynamic light scattering (DLS)**. Polydispersity of the samples was measured as intensity. SepF was used at 50 μM and FtsZ$_{CTD}$ peptide at 100 μM. SepF was monodisperse (**a**), while SUVs presented a larger polydispersity with an average diameter around 70 nm (**b**). When SepF and SUVs were incubated together, several peaks corresponding to larger particles were seen (**c**). These peaks were reversed when the FtsZ$_{CTD}$ was added to the SepF samples (**d**).

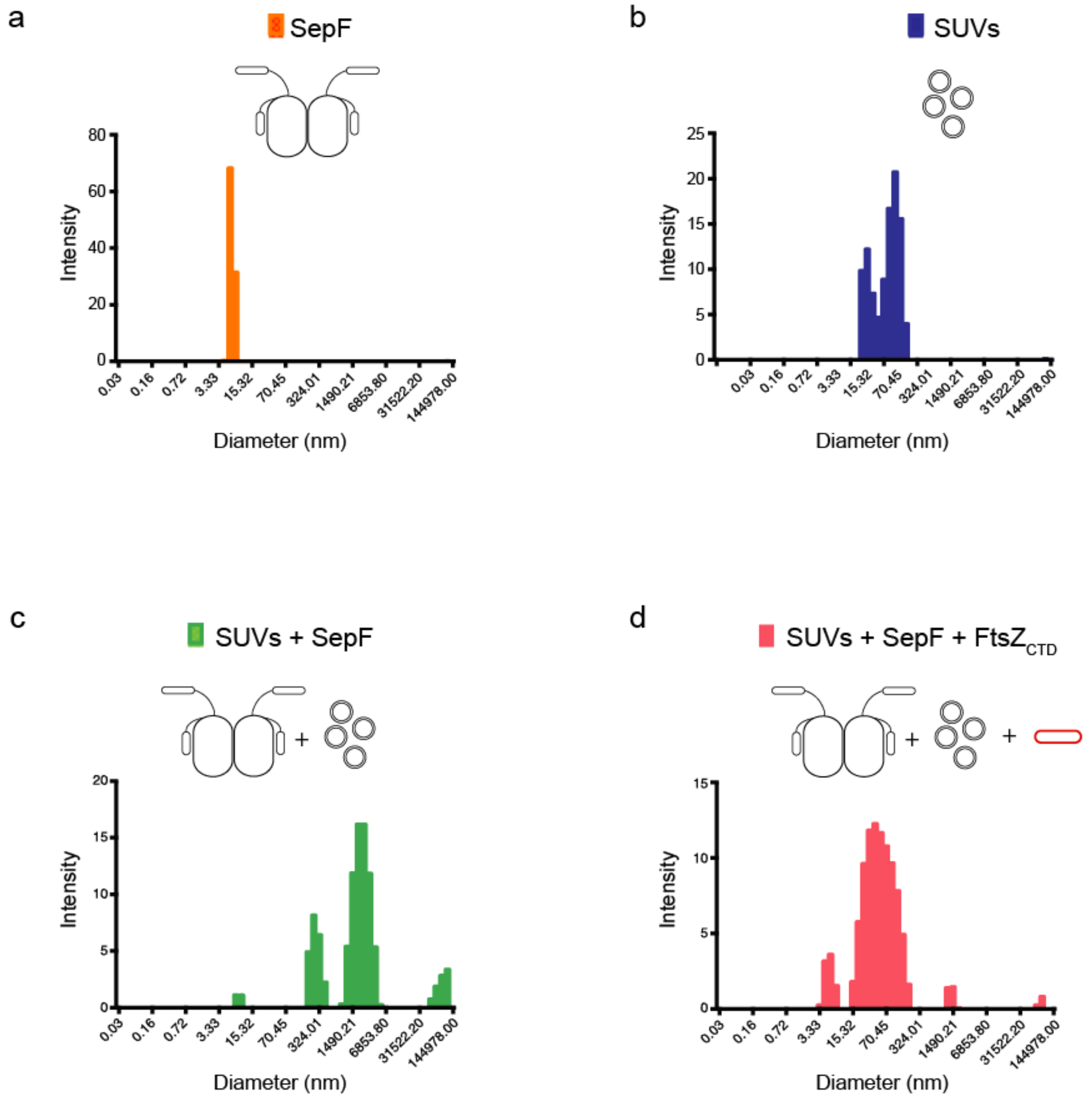

**Supplementary Figure 17: SepF$_{\Delta\alpha3}$-SUV-FtsZ$_{CTD}$ interactions. a**. In DLS, SepF$_{\Delta\alpha3}$ was monodisperse, while SUVs presented a larger polydispersity with an average diameter around 70 nm. **b.** DLS, polymerization assay and negatively-stained EM images of the end-point of the assay for SepF$_{\Delta\alpha3}$ in the presence of SUVs. **c.** Idem for SepF$_{\Delta\alpha3}$ in the presence of SUVs + FtsZ$_{CTD}$. Incubation of SUVs with SepF$_{\Delta\alpha3}$ led to formation of large particles and vesicle tubulation, but these effects were only partially reversed upon addition of FtsZ$_{CTD}$.

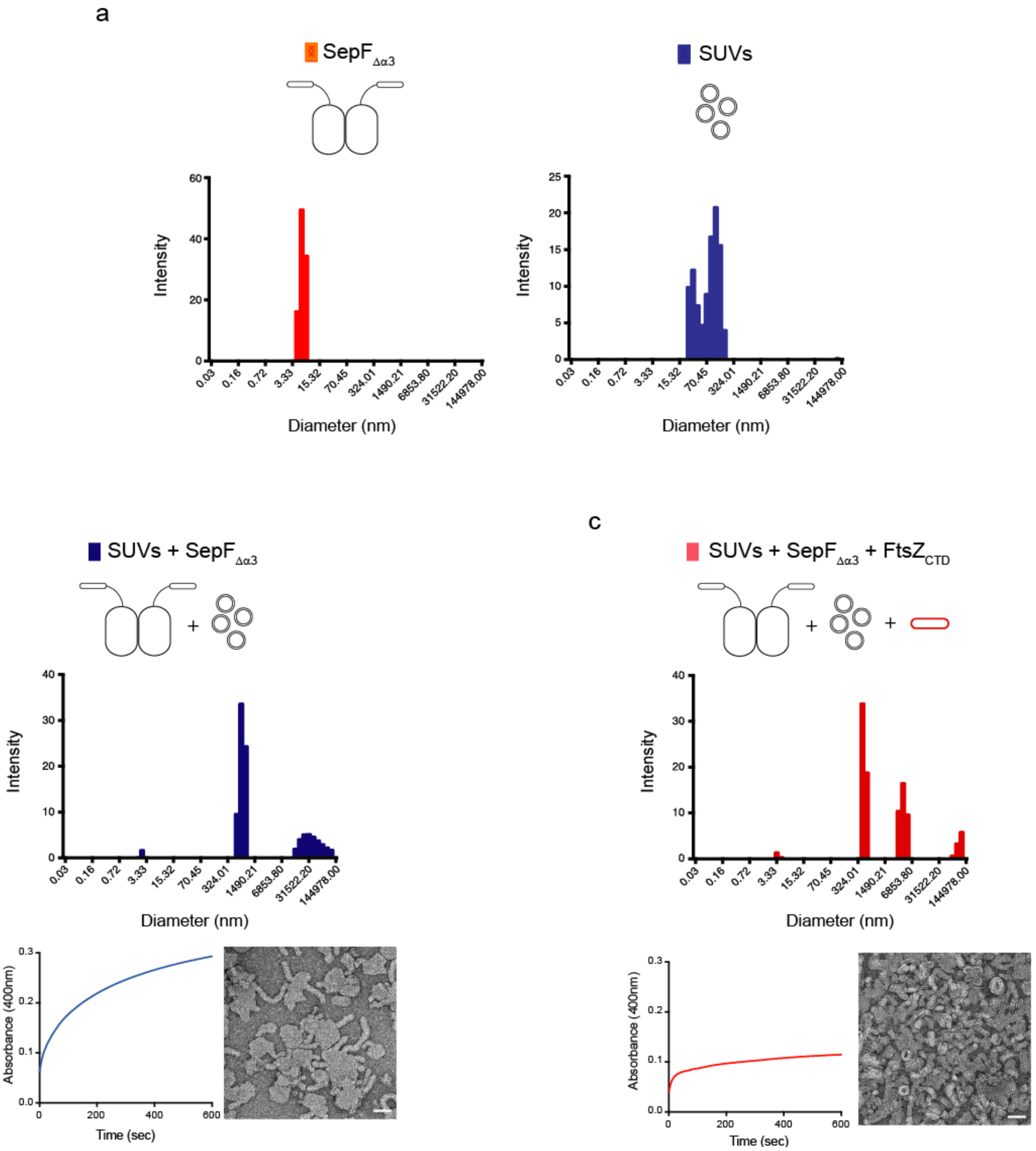

**Supplementary Figure 18: Expression and phenotypes of SepF-Scarlet and SepF$_{K125/F131A}$-Scarlet in WT *C. glutamicum*. a.** Western blot of whole cell extracts of WT-*P$_{gntK}$-sepF$_{K125/F131A}$-scarlet* (A) and WT-*P$_{gntK}$-sepF-scarlet* (B) grown either in 4% sucrose (S) or 4% sucrose + 1% gluconate (G). **b.** Violin plot of triplicate analysis showing the distribution of cell length at 5 hours of WT-*P$_{gntK}$-sepF$_{K125/F131A}$-scarlet* (purple) and WT-*P$_{gntK}$-sepF-scarlet* (blue). Medians and standard deviations of cell lengths are shown in Table T5.

a

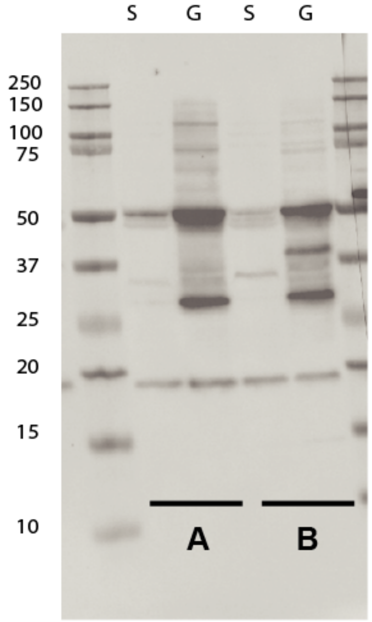

b

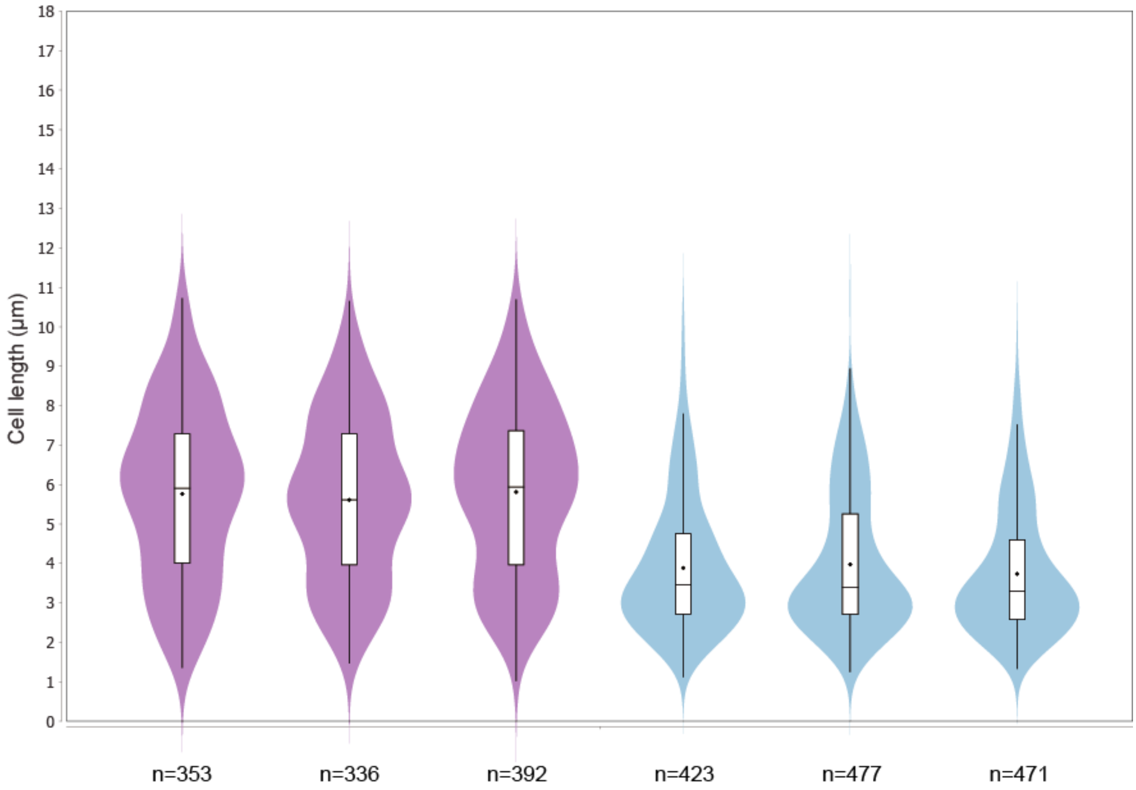

**Supplementary Figure 19: SDS-PAGE of all purified recombinant proteins used in this work**. The molecular weight markers are indicated on the left of the gel (in kDa). The protein constructs and calculated molecular weights and for each lane are shown in the figure.

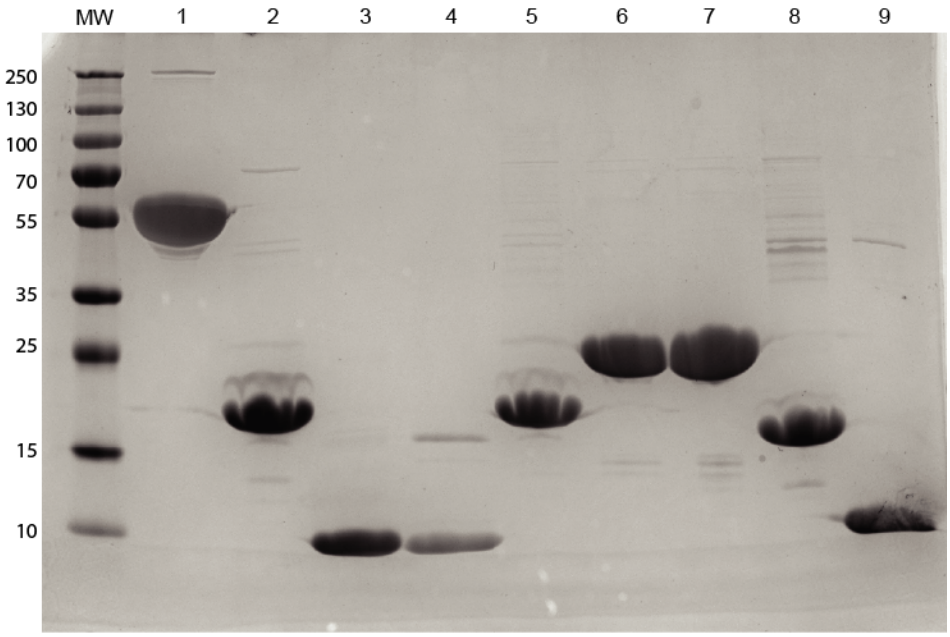

| | |
|---|---|
| 1: FtsZ | 47 kDa |
| 2: SepF | 17 kDa |
| 3: $\text{SepF}_{\Delta ML}$ | 10 kDa |
| 4: $\text{SepF}_{\Delta ML,\Delta\alpha3}$ | 8 kDa |
| 5: $\text{SepF}_{K125E,F131A}$ | 17 kDa |
| 6: His6:SUMO-$\text{SepF}_{\Delta ML,F131A}$ | 22 kDa |
| 7: His6:SUMO-$\text{SepF}_{\Delta ML,K125E,F131A}$ | 22 kDa |
| 8: $\text{SepF}_{\Delta\alpha3}$ | 15 kDa |
| 9: *M. tuberculosis* $\text{SepF}_{\Delta ML}$ | 11 kDa |

**Supplementary Figure 20: Representative electron density maps for each crystal structure of SepF determined in this work. a.** SepF$_{\Delta ML}$. **b.** SepF$_{\Delta ML}$ in complex with FtsZ$_{CTD}$. **c.** SepF$_{\Delta ML, \Delta\alpha 3}$. **d.** SepF$_{\Delta ML, \Delta\alpha 3}$ in complex with FtsZ$_{CTD}$. The (2mFobs-DFcalc) maps contoured at 1.5 $\sigma$ show the same $\alpha$-helix (residues 102-119) in all structures.

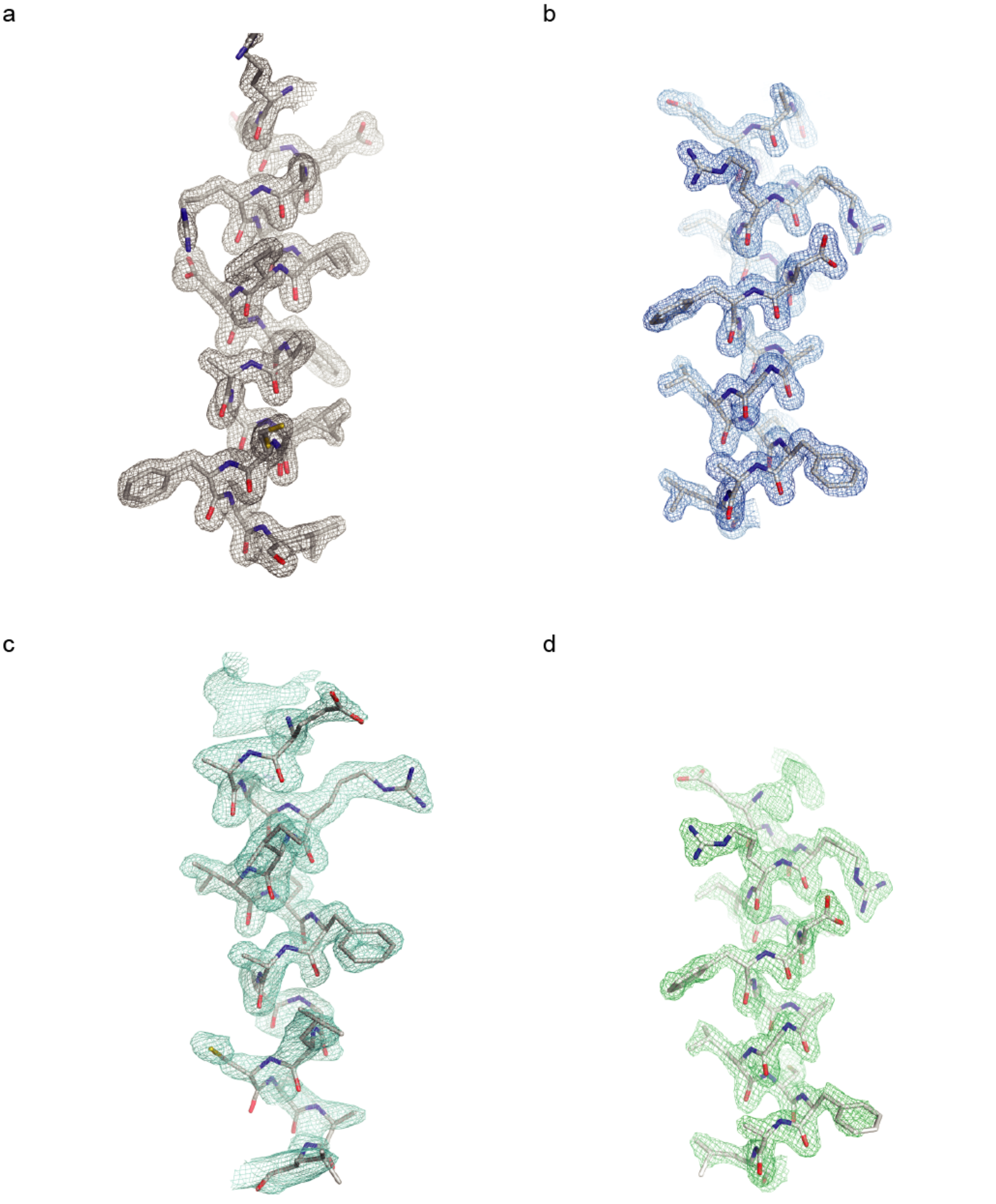

**Supplementary Figure 21: Characterization of anti-SepF and anti-FtsZ antibodies. a.** Anti-SepF (α-SepF) antibody characterization. Lanes 1-3: Serial dilutions (12.5, 6.25, 3.12 μg) of recombinant SepF against which the antibody was raised. Lanes 4-6: Serial dilutions (12.5, 6.25, 3.12 μg) of recombinant SepF$_{\Delta ML}$. Lanes 7-8: 120 μg of whole cell extracts of *C. glutamicum* WT in exponential (7) or stationary (8) phase. Lanes 9-10: 120 μg of whole cell extracts of *C. glutamicum* $P_{ino}$-*sepF* grown in 1% *myo*-inositol (SepF depletion condition) in exponential (9) or stationary (10) phase. **b.** Anti-FtsZ (α-FtsZ) antibody characterization. Lanes 1-5: Serial dilutions (100, 50, 25, 12.5, 6.25 ng) of recombinant FtsZ against which the antibody was raised. Lanes 6-7: 60 μg of whole cell extracts of *C. glutamicum* WT in exponential (6) or stationary (7) phase. Lanes 8-9: 120 μg of whole cell extracts of *C. glutamicum* WT in exponential (8) or stationary (9) phase.

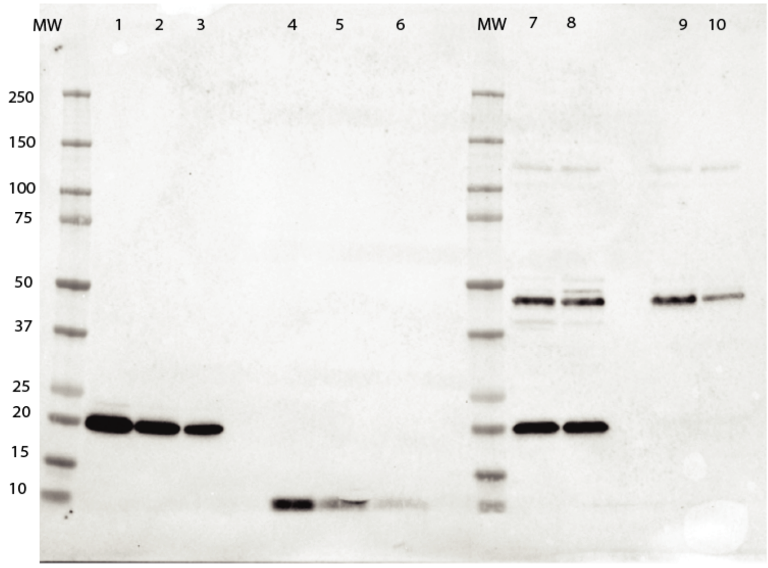

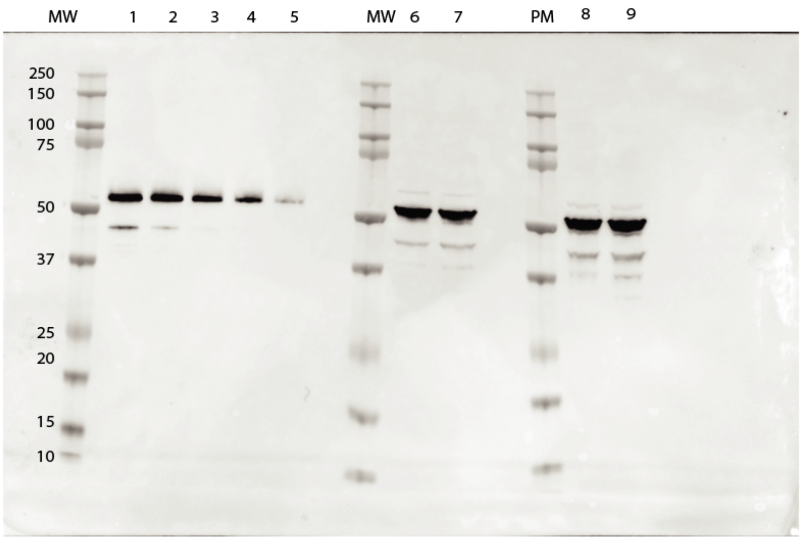

**Supplementary Figure 22: Full uncropped Western Blots of all the analyses shown in this work.** The boxes correspond to the crops used in the named figures.

Figure 1

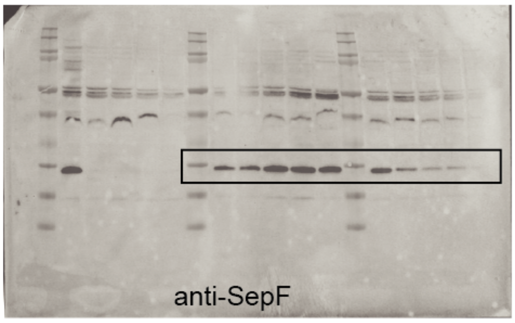

Figure 1

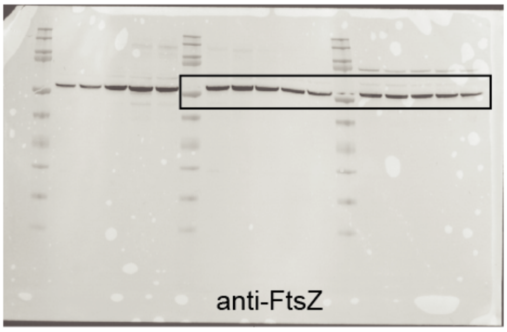

Suppl. Fig. 1

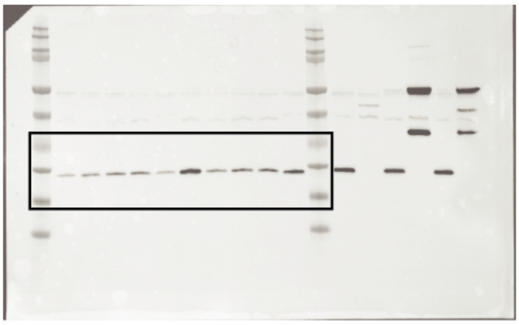

Suppl. Fig. 4

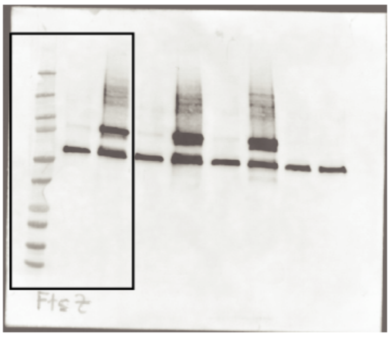

Suppl. Fig.11

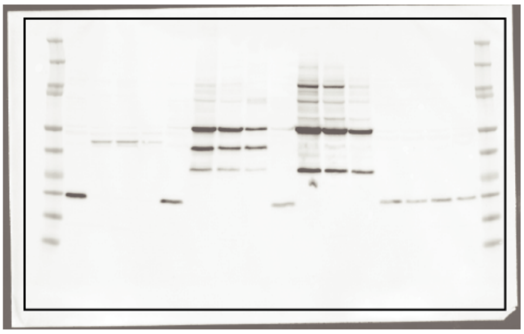

Suppl. Fig. 18

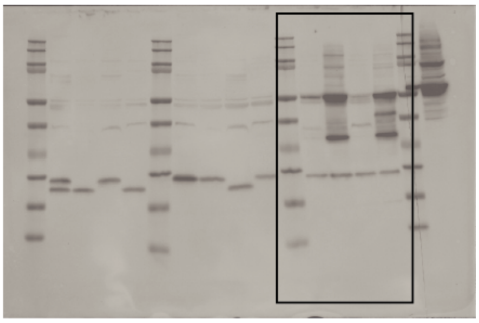

Suppl. Fig. 21

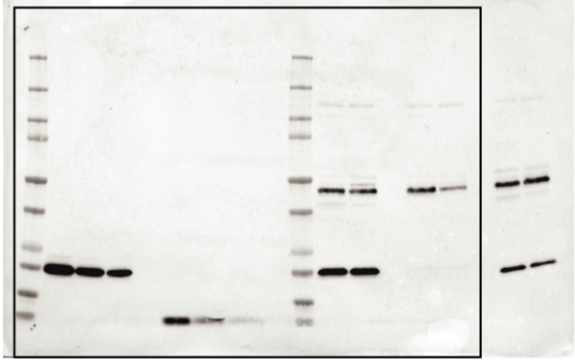

**Supplementary Table 1.** Crystallographic data.

| | **SepF$_{\Delta ML}$-FtsZ$_{CTD}$** | **SepF$_{\Delta ML}$** | **SepF$_{\Delta ML,\Delta\alpha3}$** | **SepF$_{\Delta ML,\Delta\alpha3}$-FtsZ$_{CTD}$** |
|---|---|---|---|---|
| **Data collection** | | | | |
| Space group | P $2_1$ $2_1$ $2_1$ | P $2_1$ $2_1$ $2_1$ | C 2 | P 21 |
| Cell dimensions | | | | |
| a, b, c (Å) | 33.67, 46.08, 98.98 | 35.32, 53.08, 95.63 | 65.84, 32.27, 74.41 | 34.25, 75.53, 52.61 |
| α, β, γ (°) | 90, 90, 90 | 90, 90, 90 | 90, 114.6, 90 | 90, 102.71, 90 |
| Resolution (Å)* | 49.5 – 1.6 (1.63 – 1.6) | 46.4 – 1.8 (1.84 – 1.8) | 32.9 – 1.5 (1.53 – 1.5) | 42.5 – 2.2 (2.27 – 2.2) |
| $R_{sym}$ | 0.071 (0.577) | 0.074 (0.551) | 0.026 (0.166) | 0.140 (0.555) |
| I/σ(I) | 10.3 (1.9) | 12.9 (2.5) | 19.6 (4.9) | 8.8 (3.0) |
| Completeness (%) | 96.9 (98.3) | 99.9 (100) | 95.2 (97.3) | 99.8 (99.9) |
| Redundancy | 3.6 (3.4) | 5.2 (5.2) | 2.5 (2.1) | 6.7 (6.6) |
| **Refinement** | | | | |
| Resolution (Å) | 1.6 | 1.8 | 1.50 | 2.2 |
| Number of reflections | 21381 | 17290 | 21918 | 13296 |
| R-work/R-free | 0.208 / 0.230 | 0.189 / 0.214 | 0.211 / 0.218 | 0.206 / 0.258 |
| Number of atoms | | | | |
| protein | 1439 | 1332 | 1052 | 2396 |
| ligands/ions | 0 | 6 | 0 | 1 |
| water | 148 | 109 | 84 | 115 |
| B-factors (Å$^2$) | | | | |
| protein | 26.4 | 31.3 | 39.6 | 31.6 |
| ligands/ions | - | 22.3 | - | 45.7 |
| water | 35.1 | 43.8 | 36.7 | 35.1 |
| RMS deviations | | | | |
| Bond length (Å) | 0.010 | 0.010 | 0.010 | 0.010 |
| Bond angles (°) | 0.97 | 1.02 | 1.01 | 1.09 |
| ***PDB code*** | **6SAT** | **6SCP** | **6SCQ** | **6SCS** |

* Values in parenthesis refer to the highest recorded resolution shell.

**Supplementary Table 2.** FtsZ protein levels in pull-down analyses of *C. glutamicum* strains expressing SepF-Scarlet or $SepF_{K125/F131A}$ using Scarlet as the bait protein (see Materials and Methods for details). **a**. The tables show the number of peptides and the total signal (XIC) obtained for FtsZ, Scarlet, or SepF-Scarlet constructions in three replicates of the pull-down analyses using either SepF-Scarlet or $SepF_{K125/F131A}$-Scarlet. XIC intensities were obtained using Patternlab for Proteomics XIC quantitation mode. In both cases, the Signal of SepF is estimated as Signal [SepF-Scarlet] – Signal [Scarlet] or Signal [$SepF_{K125/F131A}$-Scarlet] – Signal [Scarlet]. **b**. The table shows the average and standard deviation of the FtsZ normalized signal from IP experiments using SepF-Scarlet and $SepF_{K125/F131A}$-Scarlet. The fold change in normalized FtsZ levels between the two experiments was 7.2 (p value 0.004).

a

| | **SepF-Scarlet** | | | | | |
|---|---|---|---|---|---|---|
| | Replicate 1 | | Replicate 2 | | Replicate 3 | |
| **Signal** | **# Peptides** | **Signal (XIC)** | **# Peptides** | **Signal (XIC)** | **# Peptides** | **Signal (XIC)** |
| FtsZ | 25 | 3.94E+07 | 30 | 5.67E+07 | 24 | 4.15E+07 |
| SepF-Scarlet | 60 | 3.88E+08 | 73 | 9.85E+08 | 56 | 3.90E+08 |
| Scarlet | 38 | 2.08E+08 | 45 | 4.96E+08 | 35 | 1.95E+08 |
| SepF* | | 1.79E+08 | | 4.89E+08 | | 1.95E+08 |
| FtsZ/SepF | | 0.219 | | 0.116 | | 0.213 |

| | **SepFK125/F131A-Scarlet** | | | | | |
|---|---|---|---|---|---|---|
| | **Replicate 1** | | **Replicate 2** | | **Replicate 3** | |
| **Signal** | **# Peptides** | **Signal (XIC)** | **# Peptides** | **Signal (XIC)** | **# Peptides** | **Signal (XIC)** |
| FtsZ | 19 | 9.20E+06 | 19 | 9.89E+06 | 16 | 1.30E+07 |
| $SepF_{K125/F131A}$-Scarlet | 72 | 9.20E+08 | 63 | 5.67E+08 | 56 | 5.57E+08 |
| Scarlet | 45 | 4.07E+08 | 38 | 1.59E+08 | 31 | 1.78E+08 |
| SepF* | | 5.13E+08 | | 4.08E+08 | | 3.78E+08 |
| FtsZ/SepF | | 0.0179 | | 0.0242 | | 0.0345 |

b

| | **Signal FtsZ/SepF average** | **SD** |
|---|---|---|
| SepF-Scarlet | 0.183 | 0.058 |
| $SepF_{K125/F131A}$-Scarlet | 0.0255 | 0.0084 |

**Supplementary Table 3.** Bacterial strains and plasmids used in this study.

| Strain or plasmid | Characteristics | Reference |
|---|---|---|
| ***E. coli*** | | |
| DH5*α* | F- endA1 Φ80dlacZΔM15 Δ(lacZYA-argF)U169 recA1 relA1 hsdR17(rK–mK+) deoR supE44 thi-1 gyrA96 phoA λ–; strain used for general cloning procedures | Ref. 1 |
| BL21(DE) | F- ompT hsdSB(rB–mB–) gal dcm (DE3); host for protein production | Ref. 2 |
| CopyCutter EPI400 | F- mcrA Δ(mrr-hsdRMS-mcrBC) Φ80dlacZΔM15 ΔlacX74 recA1 endA1 araD139 Δ(ara, leu)7697 galU galK λ- rpsL (StrR) nupG trfA tonA pcnB4 dhfr | Ref. 3 |
| ***C. glutamicum*** | | |
| ATCC 13032 | Biotin-auxotrophic wild type | Ref. 4 |
| $P_{ino}$-*sepF* | *myo*-inositol dependent *sepF* silencing strain. ATCC 13032 with insertion of a terminator and $P_{ino}$ promoter to silence *sepF* (*cg2363)* expression. Repressible in the presence of *myo*-inositol | This work |
| **Plasmids** | | |
| *pK19mobsacB* | KanaR; plasmid for allelic exchange in *C. glutamicum*; (pK18 $oriV_{Ec}$, sacB, lacZα) | Ref. 5 |
| pk19-P3323-lcpA | KanaR; pK19mobsacB derivative. Used as a PCR template to amplify a transcriptional terminator and the promoter of cg3323 ($P_{ino}$) | Ref. 6 |
| *pK19-*$P_{ino}$*-sepF* | KanaR; pk19mobsacB derivative containing 500 bp upstream-region of *sepF*, a transcriptional terminator, the $P_{ino}$ promoter and 500bp of the *sepF* coding region | This work |
| *pET-SUMO-sepF* | KanaR; pET derivate for *C. glutamicum* SepF recombinant expression containing a N-terminal His-tag followed by a SUMO protease cleavage site | This work |
| *pET-SUMO-mtbsepF* | KanaR; pET derivate for *M. tuberculosis* SepF recombinant expression containing a N-terminal His-tag followed by a SUMO protease cleavage site | This work |
| *pET-SUMO-ftsZ* | KanaR; pET derivate for *C. glutamicum* FtsZ recombinant expression containing an N-terminal His-tag followed by a SUMO protease cleavage site | This work |
| *pET-SUMO-mtbsepF*$_{\Delta ML}$ | KanaR; pET derivate for *M. tuberculosis* SepF (122-218) recombinant expression containing a N-terminal His-tag followed by a SUMO protease cleavage site | This work |
| *pET-SUMO-sepF*$_{\Delta ML}$ | KanaR; pET derivate for *C. glutamicum* SepF (63-152) recombinant expression containing a N-terminal His-tag followed by a SUMO protease cleavage site | This work |
| *pET-SUMO-sepF*$_{\Delta ML,\Delta\alpha3}$ | KanaR; pET derivate for *C. glutamicum* SepF (63-137) recombinant expression containing a N-terminal His-tag followed by a SUMO protease cleavage site | This work |
| *pET-SUMO-sepF*$_{\Delta\alpha3}$ | KanaR; pET derivate for *C. glutamicum* SepF (1-137) recombinant expression containing a N-terminal His-tag followed by a SUMO protease cleavage site | This work |
| *pET-SUMO-sepF*$_{\Delta ML,F131A}$ | KanaR; pET derivate for *C. glutamicum* SepF (63-152) recombinant expression containing a N-terminal His-tag followed by a SUMO protease cleavage site. Carrying amino acid substitution F131A | This work |
| *pET-SUMO-sepF*$_{\Delta ML,K125/F131A}$ | KanaR; pET derivate for *C. glutamicum* SepF (63-152) recombinant expression containing a N-terminal His-tag followed by a SUMO protease cleavage site. Carrying double amino acid substitution K125E and F131A | This work |

| | | |
|---|---|---|
| *pET-SUMO-sepF$_{K125/F131A}$* | KanaR; pET derivate for *C. glutamicum* SepF recombinant expression containing a N-terminal His-tag followed by a SUMO protease cleavage site. Carrying double amino acid substitution K125E and F131A | This work |
| *pTGR5* | KanaR; *E. coli/C. glutamicum* shuttle vector for regulated gene expression under control of tac promoter ($P_{tac}$ lacI ColE1 $oriV_{Ec}$ pGA1 $oriV_{Cg}$) | Ref. 7 |
| *pCLTON1* | KanaR; *E. coli* / *C. glutamicum* shuttle expression vector with the *B. subtilis* derived $P_{tet}$ promoter from pWH105 and the tetR gene under control of *C. glutamicum* $P_{gap}$ promoter from pJC1-pgap-tetR. | Ref. 8 |
| *pMA-mscarlet-I* | AmpR; pMA vector (GeneArt, Thermo Fisher Scientific) containing a synthetic gene coding for mScarlet-I, codon optimized for expression in *C. glutamicum* | This work |
| *pMA-mneongreen* | AmpR; pMA vector (GeneArt, Thermo Fisher Scientific) containing a synthetic gene coding for mNeonGreen, codon optimized for expression in *C. glutamicum* | This work |
| *pUMS3* | KanaR; pTGR5 derivative in which $P_{tac}$ was exchanged by $P_{gntK}$ promoter | This work |
| *pUMS3-scarlet* | KanaR; pUMS3 derivative for expression of Scarlet under control of $P_{gntK}$ promoter | This work |
| *pUMS3-sepF-scarlet* | KanaR; pUMS3 derivative for expression of cgSepF-Scarlet under control of $P_{gntK}$ promoter | This work |
| *pUMS3-sepF$_{K125/F131A}$-scarlet* | KanaR; pUMS3 derivative for expression of cgSepF$_{K125/F131A}$-Scarlet under control of $P_{gntK}$ promoter | This work |
| *pUMS3-sepF$_{\Delta ML}$-scarlet* | KanaR; pUMS3 derivative for expression of cgSepF$_{\Delta ML}$-Scarlet under control of $P_{gntK}$ promoter | This work |
| *pUMS3-mneon-ftsZ* | KanaR; pUMS3 derivative for expression of mNeon-cgFtsZ under control of $P_{gntK}$ promoter | This work |
| *pUMS40* | KanaR; pTGR5 derivative in which $P_{tac}$ was exchanged by $P_{tet}$ promoter | This work |
| *pUMS40-sepF* | KanaR; pUMS40 derivative for expression of cgSepF under control of $P_{tet}$ promoter | This work |
| *pUMS40-sepF$_{\Delta\alpha3}$* | KanaR; pUMS40 derivative for expression of cgSepF$_{\Delta\alpha3}$ under control of $P_{tet}$ promoter | This work |
| *pUMS40-sepF-scarlet* | KanaR; pUMS40 derivative for expression of cgSepF-Scarlet under control of $P_{tet}$ promoter | This work |

**Supplementary Table 4:** Oligonucleotides used in this study

| Oligonucleotide | Sequence (5' → 3') and properties[a] |
|---|---|
| **Construction of pET-SUMO-cgSepF$_{\Delta ML}$** | |
| P1 | AGTTATCAGTCCACCATTGTTCCGGTT |
| P2 | GGTGGACTGATAACTCATGCCACCAATCTG |
| **Construction of pET-SUMO-cgSepF$_{\Delta ML,\Delta\alpha 3}$ & pET-SUMO-cgSepF$_{\Delta\alpha 3}$** | |
| P13 | GGAATAGTCTAACATTAGCACGTCT |
| P14 | AGACGTGCTAATGTTAGACTATTCC |
| **Construction of pET-SUMO-cgSepF$_{\Delta ML,F131A}$** | |
| P39 | TTACCGCCGCCGTCGTG |
| P40 | GATAGCGTTACCGCCGCCGT |
| **Construction of pET-SUMO-cgSepF$_{\Delta ML,K125/F131A}$** | |
| P33 | TAAAATGCAGGAAATCGATAGCG |
| P34 | AACGCTATCGATTTCCTGCATTTTAC |
| **Construction of pET-SUMO-mtbSepF$_{\Delta ML}$** | |
| P7 | GATGGTCACCCGC |
| P8 | CGGGTGACCATCGCCACCAATCTGTTC |
| **Construction of pk19-*P$_{ino}$-sepF* and the respective mutant strain** | |
| P123 | **TGAAGGGAAACTGCC**ATAAAACGAAAGGCTCAGTCGAAAGAC |
| P124 | **CTTGAGCATGGACAT**CTAAAATTTCTCCTCTTAAAAAGATAACGGCC |
| P121 | **TGTTGTGTGGAATTG**TACAAGAGTTTGGTGTCACCGC |
| P122 | GGCAGTTTCCCTTCACCTG |
| P125 | ATGTCCATGCTCAAGAAGACTAAAGAA |
| P126 | **AATTGTTATCCGCTCA**AGACCTTGGAAAAGAGCCGTAGAGG |
| **Primers for colony PCR used during the construction of the *P$_{ino}$-sepF* strain** | |
| P86 | CTTGACTGCAGCCATGGTTT |
| P70 | TGTTGTGTGGAATTGCCAAAGCACTCGAACTTCC |
| P142 | CTCATCGCTGTAGTGACCCGGTA |
| P85 | GATTACCGTCGTGATGAGCG |
| **Construction of pUMS3** | |
| OligoMM_157 | CTCACGCACTCCGGGTATCTAGATGACATACGAACAAATCGTTGATCTAGTT |
| OligoMM_158 | GTTCGGTGAGGTCATATGGTCTTATCCTTTCTTTGGTGGCGTCTC |
| **Construction of pUMS3-cgSepF-Scarlet** | |
| OligoMM_178 | **ATGGTCTTATCCTTTCTTTG**GTGGC |
| OligoMM_179 | **AGCGGCCGCTTAAGGTAC** |
| OligoMM_183 | **CAAAGAAAGGATAAGACCAT**ATGTCCATGCTCAAGAAGACTAAAGAATTCTTCGGACTCG |
| OligoMM_184 | **GCCAGATCCCTCGAGGCGGATG**CGTGCGGCGCG |
| OligoMM_180 | **ACGCATCCGCCTCGAGGGATCTGGC**CAGGGACCGGGCTCAGGCCAAGGAAGCGGCCATATGGTGTCCAAGGGCGAAG |
| OligoMM_181 | **CGGTACCTTAAGCGGCCGCT**TTACTTGTACAGTTCATCCATGCC |
| **Construction of pUMS3-cgSepF$_{K125/F131A}$-Scarlet** | |
| OligoMM_188 | AGATTGACAGCGTCACCGCCGCTGTCGTTCCAGAGCTGTCCAACATCAGCAC |
| OligoMM_189 | GCGGTGACGCTGTCAATCTCCTGCATCTTGCCACGCAATGCGAAGCACAG |
| **Construction of pUMS3-cgSepF$_{\Delta ML}$-Scarlet** | |
| OligoMM_178 | **ATGGTCTTATCCTTTCTTTG**GTGGC |
| OligoMM_179 | **AGCGGCCGCTTAAGGTAC** |
| OligoMM_187 | **CAAAGAAAGGATAAGACCAT**ATGTCTTACCAGTCCACCATCGTTCCAGTAGAGCTTCATTC |

| | |
|---|---|
| OligoMM_184 | **GCCAGATCCCTCGAGGCGGATGCGTG**CGGCGCG |
| OligoMM_180 | **ACGCATCCGCCTCGAGGGATCTGGC**CAGGGACCGGGCTCAGGCCAAGGAAGCGGCCATATGGTGT CCAAGGGCGAAG |
| OligoMM_181 | **CGGTACCTTAAGCGGCCGCT**TTACTTGTACAGTTCATCCATGCC |
| **Construction of pUMS3-mNeon-cgFtsZ** | |
| OligoMM_178 | **ATGGTCTTATCCTTTCTTTGGT**GGC |
| OligoMM_179 | **AGCGGCCGCTTAAGGTAC** |
| OligoMM_190 | **CAAAGAAAGGATAAGACCAT**ATGGTGTCCAAGGGCGAAG |
| OligoMM_191 | **GCCAGATCCCTCGAGCTTGTACAGT**TCATCCATGCCC |
| OligoMM_192 | **ACTGTACAAGCTCGAGGGATCTGGC**CAGGGACCGGGCTCAGGCCAAGGAAGCGGCATGACCTCAC CGAACAAC |
| OligoMM_193 | **CGGTACCTTAAGCGGCCGCT**TTACTGGAGGAAGCTGGG |
| **Construction of pUMS40** | |
| OligoMM_254 | **AATATGCGGCCGCATATATG**GATC |
| OligoMM_227 | **ATGGTGAGCAAGGGCGAGGAG** |
| OligoMM_252 | **CATATATGCGGCCGCATATT**CCTGCCGCCAGCGGGCGTACAAAAGTG |
| OligoMM_253 | **TGAACAGCTCCTCGCCCTTGCTCACCAT**AGTGTATCAACAAGCTGGGGATCTTAAGCTTG |
| **Construction of pUMS40-cgSepF** | |
| OligoMM_202 | ATCCGCTAACTCGAGGGATCTGGCCAGGGACCGGG |
| OligoMM_203 | CTCGAGTTAGCGGATGCGTGCGGCGCGCTCGAGC |
| **Construction of pUMS40-cgSepF$_{\Delta\alpha3}$** | |
| OligoMM_204 | CCAGAGTAACTCGAGGGATCTGGCCAGGGACC |
| OligoMM_205 | CTCGAGTTACTCTGGAACGACAGCGAAGGTGACGC |
| **Construction of pUMS40-cgSepF-Scarlet** | |
| OligoMM_258 | **AGTGTATCAACAAGCTGGGGA** |
| OligoMM_179 | **AGCGGCCGCTTAAGGTAC** |
| OligoMM_259 | **TCCCCAGCTTGTTGATACACT**ATGTCCATGCTCAAGAAGACTAAAGAATTCTTCGGACTCG |
| OligoMM_184 | **GCCAGATCCCTCGAGGCGGATGCGT**GCGGCGCG |
| OligoMM_180 | **ACGCATCCGCCTCGAGGGATCTGGC**CAGGGACCGGGCTCAGGCCAAGGAAGCGGCCATATGGTGT CCAAGGGCGAAG |
| OligoMM_181 | **CGGTACCTTAAGCGGCCGCT**TTACTTGTACAGTTCATCCATGCC |

[a] Overlaps for Gibson assembly are written in bold letters. Restriction sites are underlined.

**Supplementary Table 5.** Statistical analysis of cell lengths in all violin plots.

| Strain + construct | Time point | Replicate | N cells | Mean length | Standard dev. |
|---|---|---|---|---|---|
| **Fig. 1** | | | | | |
| WT | t = 0 | 1 | 348 | 3,10 | 0,63 |
| | t = 3 | 1 | 362 | 3,30 | 0,71 |
| | t = 6 | 1 | 314 | 2,63 | 0,61 |
| $P_{ino}$-*sepF* | t = 0 | 3 | 413 | 2,40 | 0,49 |
| | t = 3 | 3 | 339 | 4,11 | 0,90 |
| | t = 6 | 3 | 318 | 7,19 | 1,56 |
| **Fig. 3** | | | | | |
| $P_{ino}$-*sepF* - $P_{gntK}$-$sepF_{\Delta ML}$-*scarlet* | t = 6 | 1 | 302 | 9,81 | 2,24 |
| $P_{ino}$-*sepF* - $P_{gntK}$-*sepF-scarlet* | t = 6 | 1 | 379 | 5,38 | 1,62 |
| $P_{ino}$-*sepF* - $P_{gntK}$-$sepF_{K125/F131A}$-*scarlet* | t = 6 | 1 | 328 | 9,96 | 2,85 |
| $P_{ino}$-*sepF* - empty vector | t = 6 | 1 | 466 | 2,61 | 0,57 |
| **Supplementary Fig. 1** | | | | | |
| WT - empty vector | t = 4.5 | 1 | 369 | 2,61 | 0,56 |
| | | 2 | 329 | 2,58 | 0,57 |
| | | 3 | 372 | 2,57 | 0,58 |
| $P_{ino}$-*sepF* - $P_{tet}$-*sepF* | t = 4.5 | 1 | 351 | 2,61 | 0,58 |
| | | 2 | 406 | 2,61 | 0,54 |
| | | 3 | 436 | 2,68 | 0,57 |
| **Supplementary Fig. 2** | | | | | |
| WT | t = 0 | 1 | 348 | 3,10 | 0,63 |
| | | 2 | 413 | 3,04 | 0,65 |
| | | 3 | 337 | 3,11 | 0,68 |
| | t = 3 | 1 | 362 | 3,30 | 0,71 |
| | | 2 | 293 | 3,18 | 0,72 |
| | | 3 | 338 | 3,20 | 0,69 |
| | t = 6 | 1 | 314 | 2,63 | 0,61 |
| | | 2 | 424 | 2,55 | 0,60 |
| | | 3 | 365 | 2,61 | 0,58 |
| $P_{ino}$-*sepF* | t = 0 | 1 | 349 | 2,54 | 0,51 |
| | | 2 | 298 | 2,66 | 0,51 |
| | | 3 | 413 | 2,40 | 0,49 |
| | t = 3 | 1 | 242 | 4,30 | 0,89 |
| | | 2 | 321 | 4,19 | 1,00 |

| | | | | | |
|---|---|---|---|---|---|
| | | 3 | 339 | 4,11 | 0,90 |
| | t = 6 | 1 | 307 | 7,71 | 1,44 |
| | | 2 | 314 | 7,60 | 1,84 |
| | | 3 | 318 | 7,19 | 1,56 |
| **Supplementary Fig. 4** | | | | | |
| WT - *P<sub>gntK</sub>-mneon-ftsZ* | t = 3 | 1 | 383 | 2,52 | 0,56 |
| | | 2 | 398 | 2,44 | 0,56 |
| | | 3 | 429 | 2,51 | 0,53 |
| WT - empty vector | t = 4.5 | 1 | 369 | 2,61 | 0,56 |
| | | 2 | 329 | 2,58 | 0,57 |
| | | 3 | 372 | 2,57 | 0,58 |
| **Supplementary Fig. 5** | | | | | |
| $P_{ino}$-*sepF* - $P_{gntK}$-*mneon-ftsZ* | t = 0 | 1 | 251 | 2,85 | 0,64 |
| | | 2 | 398 | 2,99 | 0,64 |
| | | 3 | 337 | 2,88 | 0,71 |
| | t = 3 | 1 | 326 | 5,59 | 1,37 |
| | | 2 | 350 | 5,59 | 1,31 |
| | | 3 | 317 | 5,74 | 1,17 |
| | t = 6 | 1 | 295 | 9,98 | 2,31 |
| | | 2 | 352 | 9,99 | 1,98 |
| | | 3 | 297 | 9,90 | 1,83 |
| **Supplementary Fig. 11** | | | | | |
| $P_{ino}$-*sepF* - $P_{gntK}$-*sepF*$_{\Delta ML}$-*scarlet* | t = 6 | 1 | 302 | 9,81 | 2,24 |
| | | 2 | 313 | 9,33 | 1,97 |
| | | 3 | 377 | 9,46 | 2,07 |
| $P_{ino}$-*sepF* - $P_{gntK}$-*sepF-scarlet* | t = 6 | 1 | 379 | 5,38 | 1,62 |
| | | 2 | 345 | 4,97 | 1,63 |
| | | 3 | 360 | 4,60 | 1,64 |
| $P_{ino}$-*sepF* - $P_{gntK}$-*sepF*$_{K125/F131A}$-*scarlet* | t = 6 | 1 | 328 | 9,96 | 2,85 |
| | | 2 | 324 | 9,74 | 2,24 |
| | | 3 | 342 | 9,52 | 2,20 |
| $P_{ino}$-*sepF* - empty vector | t = 6 | 1 | 466 | 2,61 | 0,57 |
| | | 2 | 375 | 2,52 | 0,55 |
| | | 3 | 359 | 2,58 | 0,61 |
| **Supplementary Fig. 18** | | | | | |
| WT - $P_{gntK}$-*sepF*$_{K125/F131A}$-*scarlet* | t = 5 | 1 | 353 | 5,76 | 2,18 |
| | | 2 | 336 | 5,61 | 2,05 |
| | | 3 | 392 | 5,81 | 2,13 |

| | | | | | |
|---|---|---|---|---|---|
| WT - $P_{gntK}$-*sepF-scarlet* | t = 5 | 1 | 423 | 3,88 | 1,66 |
| | | 2 | 477 | 3,97 | 1,73 |
| | | 3 | 471 | 3,73 | 1,58 |
